# Heating quinoa shoots results in yield loss by inhibiting fruit production and delaying maturity

**DOI:** 10.1101/727545

**Authors:** Jose C. Tovar, Carlos Quillatupa, Steven T. Callen, S. Elizabeth Castillo, Paige Pearson, Anastasia Shamin, Haley Schuhl, Noah Fahlgren, Malia A. Gehan

**Affiliations:** Donald Danforth Plant Science Center, St. Louis, MO, USA; Bayer U.S. – Crop Science, St. Louis, MO, USA

**Keywords:** Heat, yield, quinoa, fruit production, plant maturity, phenomics, RNA-seq

## Abstract

Increasing global temperatures and a growing world population create the need to develop crop varieties that yield more in warmer climates. There is growing interest in expanding quinoa cultivation, because of quinoa’s ability to produce nutritious grain in poor soils, with little water and at high salinity. However, the main limitation to expanding quinoa cultivation is quinoa’s susceptibility to temperatures above ~32°C. This study investigates the phenotypes, genes, and mechanisms that may affect quinoa seed yield at high temperatures. By using a differential heating system where only roots or only shoots were heated, quinoa yield losses were attributed to shoot heating. Plants with heated shoots lost 60% to 85% yield as compared to control. Yield losses were due to lower fruit production, which lowered the number of seeds produced per plant. Further, plants with heated shoots had delayed maturity and more non-reproductive shoot biomass, while plants with both heated roots and heated shoots produced more yield from panicles that escaped heat than control. This suggests that quinoa uses a type of avoidance strategy to survive heat. Gene expression analysis identified transcription factors differentially expressed in plants with heated shoots and low yield that had been previously associated with flower development and flower opening. Interestingly, in plants with heated shoots, flowers stayed closed during the day while control flowers were open. Although a closed flower may protect floral structures, this could also cause yield losses by limiting pollen dispersal, which is necessary to produce fruit in quinoa’s mostly female flowers.

**Significance Statement:** This study provides evidence that heating quinoa during flowering results in seed yield loss by lowering fruit production. Plants with low yield after heat treatment also matured more slowly, suggesting that quinoa may use a type of avoidance strategy to survive heat stress conditions. Genes differentially expressed under heat include genes involved in flower development and flower opening.

## Introduction

Global temperatures are estimated to increase 1°C to 5°C (Callery *et al.*, 2018), while the world population will grow by ~47% in the 21st century (United Nations, 2017). On average, more than half of human caloric intake comes directly from grain consumption (Awika, 2011), and feed used for meat production is 28% grain (Herrero *et al.*, 2013). Thus, there is a general need to increase grain production in a warming environment. Although farming technology, farm land expansion, and breeding have increased absolute grain yields, percent yield gains for grain crops have been decreasing in recent years (Grassini *et al.*, 2013). Grain crops are predicted to lose 3.1% to 7.4% yield for every 1°C increase unless new varieties are developed for warmer temperatures (Zhao *et al.*, 2017). Climate change-induced increases in temperature and reductions in precipitation have already caused yield losses of up to 5.5% from grain crops between 1980 and 2008 (Lobell *et al.*, 2011), a period where temperatures increased only ~0.6°C (Hansen *et al.*, 2010). To meet future global grain demands, it is vital to develop grain crop varieties adapted to higher temperatures (Challinor *et al.*, 2014) and gain a better understanding of the molecular mechanisms that influence losses in grain yield.

Quinoa (*Chenopodium quinoa* Willd.) is a grain crop (pseudocereal) grown in areas with average temperatures of 9°C to 30°C (Bhargava *et al.*, 2007a). There is growing interest in expanding quinoa cultivation (Choukr-Allah *et al.*, 2016; Jacobsen, 2003; Bazile *et al.*, 2016; Maliro *et al.*, 2017; Pulvento *et al.*, 2010), because of its nutritious grain (Vega-Gálvez *et al.*, 2010; Choukr-Allah *et al.*, 2016; Repo-Carrasco *et al.*, 2003), and its ability to grow on poor soils (Jacobsen *et al.*, 2003). However, heat is a major limitation to expanding quinoa cultivation (Hinojosa, Matanguihan, *et al.*, 2018; Lesjak and Calderini, 2017), and quinoa generally does poorly in climates with average temperatures higher than 32°C (Hinojosa, Matanguihan, *et al.*, 2018; Bazile *et al.*, 2016). A goal of this study is to gain a better understanding of how heat limits grain production in quinoa.

Quinoa’s nutritious grain and ability to grow on poor soils suggests that quinoa could be an interesting crop to study nutrient uptake. We are interested in how heat stress affects processes of nutrient uptake in the roots and grain production in the shoots, since there are few studies on how roots and shoots differentially respond to heat stress. Previous studies have shown that plants respond differently to heat in the roots compared to heat in the shoots (Heckathorn *et al.*, 2013). In wheat, root heating had a more pronounced effect than shoot heating on grain yield, shoot biomass, and root biomass (Kuroyanagi and Paulsen, 1988). In the grass, *Agrostis palustris*, photosynthesis was more severely affected when heating roots than when heating shoots (Xu and Huang, 2000; Huang *et al.*, 2001). These studies suggest that root and shoot responses to heat may involve different mechanisms. Thus, studying how quinoa roots and shoots respond to heat can provide important insights into the mechanisms involved in yield losses.

This study investigates quinoa’s phenotypic and transcriptomic responses to root heating compared to shoot heating in the sequenced accession QQ74 (PI 614886, (Jarvis *et al.*, 2017). The impact of heat on plant development, yield, and gene expression are examined in detail and we propose a possible mechanism for yield loss under heat stress.

## Results and discussion

### Shoot heating significantly decreases yield, but root heating does not have a significant effect on yield

To study the responses of quinoa to root and shoot heat stress, a sandbox system was designed and built that allowed for independent temperature control of quinoa roots and shoots (Figure S1). Since previous studies have indicated that flowering is the most susceptible developmental stage to heat stress (Lesjak and Calderini, 2017), we heat-treated quinoa during the flowering stage (~ 35 to 40 days old plants). Heat was applied during flowering since previous studies found that flowering was the most susceptible developmental stage to heat (Lesjak and Calderini, 2017). We found that soil temperature was 30°C when the air temperature was 35°C, therefore 30°C was selected as a heat treatment for roots. In total there were four treatments: 1) control treatment with roots and shoots at 22°C; 2) heated roots (HR), with roots at 30°C and shoots at 22°C; 3) heated shoots (HS), with shoots at 35°C and roots at 22°C and 4) heated roots and shoots (HRS), with roots at 30°C and shoots at 35°C. Flowering plants were heat treated for 11 days and then returned to control temperature conditions until harvest. For more detail on the sandbox set-up please see the Experimental Procedures section on “Plant material and growth conditions”.

Data analysis was done using a factorial design, to quantify the effects of root versus shoot heating. A treatment by treatment comparison was done to identify pairwise differences. For more details on the methods used for data analysis please see the Experimental Procedures section on “Statistical analysis”.

In this study yield was defined as grams of seed produced per plant. Shoot heating resulted in significant seed yield loss (two-way ANOVA on the effect of shoot heating, p-value < 0.0001). HS plants produced an average of 68% less seed yield relative to control plants (Wilcoxon rank sum test, p-value = 0.0043), and HRS plants had 61% less seed yield than control plants (Wilcoxon rank sum test, p-value = 0.0043, Figure 1a). Interestingly, root heating did not have a significant effect on seed yield (two-way ANOVA on the effect of root heating, p-value = 0.9658), with HR plants showing no significant difference from control plants (Wilcoxon rank sum test, p-value = 0.938). An experimental replicate of total plant yield showed similar yield losses, with significantly lower seed yield compared to control (two-way ANOVA on the effect of shoot heating, p-value < 0.0001) and with no significant effect from root heating (two-way ANOVA on the effect of root heating, p-value = 0.0856). In the second experimental replicate, plants with HS had an average of 85% less yield than control plants (Wilcoxon rank sum test, p-value = 0.000021), and plants with HRS produced an average of 81% less yield than control plants (Wilcoxon rank sum test, p-value = 0.000021). In contrast, plants with HR did not have a significantly lower yield than control plants (Wilcoxon rank sum test, p-value = 0.2957, Figure 1b). Overall, these results indicate that heating quinoa shoots results in greater yield losses than heating roots.

**Figure 1.**
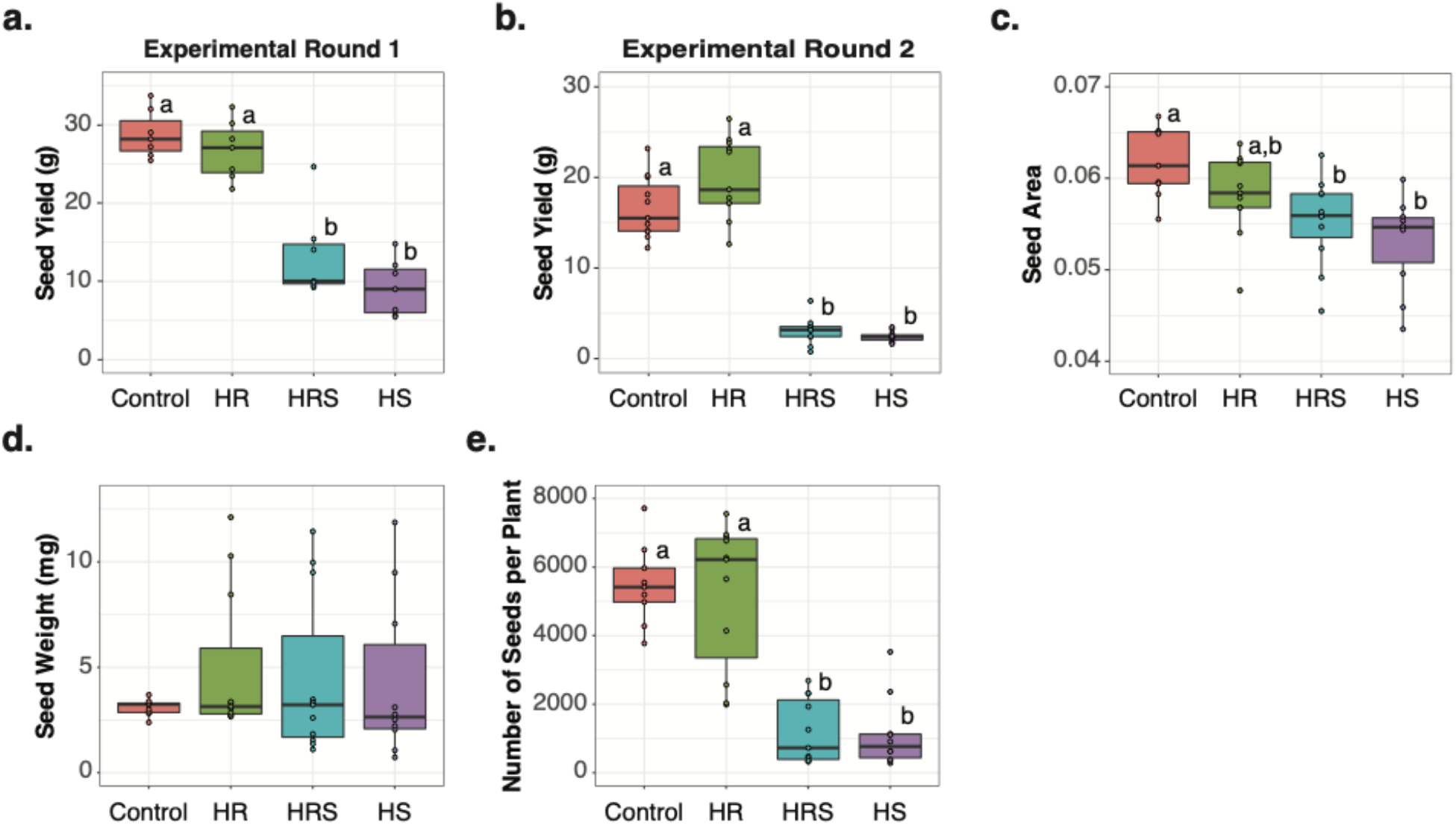
Yield analysis. a) Experimental round 1 yield per plant for each treatment (n=15 plants). b) Experimental round 2 yield per plant for each treatment (n=15 plants). c) Average normalized seed area per plant for each treatment (n=15 plants). d) Average estimated individual seed weight per plant for each treatment (n=15 plants). e) Estimated number of seeds produced per plant for each treatment (n=15 plants). Letters above boxes represent statistical significance at p-value < 0.05 from a Wilcoxon rank sum test.

Although no other quinoa studies have looked at the differential effects of heat on quinoa roots and shoots, previous studies on quinoa have found varied yield responses to heat. Quinoa accession Regalona lost 31% yield while accession BO5 lost 23% yield when night temperatures were 22°C instead of 18°C (control) during flowering (Lesjak and Calderini, 2017; Hinojosa, González, *et al.*, 2018). Hinojosa et al. 2018 found that quinoa accessions QQ74 (used in this study) and 17GR did not lose any yield when treated at 40°C day/24°C night during flowering, as compared to control at 22°C day/16°C night (Hinojosa, Matanguihan, *et al.*, 2018). The differences in QQ74 yield under heat in this study compared to Hinojosa et al. 2018 (Hinojosa, Matanguihan, *et al.*, 2018), may be due to the methodology used for treatment, including the temperatures used, and the duration of treatment. One similarity between studies that found yield losses under heat stress (this study; (Lesjak and Calderini, 2017; Hinojosa, González, *et al.*, 2018)) was a high night-time temperature treatment during flowering. Similarly, in wheat and rice, a high temperature treatment at night has been shown to negatively impact yield (Shi *et al.*, 2016; Narayanan *et al.*, 2015; Shi *et al.*, 2013; Jagadish *et al.*, 2015).

### Yield losses from shoot heating are mainly due to a smaller number of seeds produced

To investigate the changes in overall seed yield, we examined if yield losses were largely due to changes in seed size or seed number. We analyzed seed size by two independent measurements: 1) average seed weight and 2) average seed area (measured from images). Seed number was estimated by dividing the total seed weight by the average seed weight for each plant. For more information on how seed size and seed number were measured, please see the “Main panicle, seed and whole plant imaging” section in Experimental Procedures.

Shoot heating had a significant effect on seed area (two-way ANOVA on the effect of shoot heating, p-value = 0.0455), while root heating did not (two-way ANOVA on the effect of root heating, p-value = 0.708). HS plants had 14% smaller seed area than control plants (Wilcoxon rank sum test, p-value = 0.0095, Figure 1c) and HRS plants had 11% smaller seed area than control plants (Wilcoxon rank sum test, p-value = 0.0312, Figure 1c). Similarly, Bertero et al. 1999 reported a 14% smaller seed diameter with heat treatment in quinoa accession Kancolla (Bertero *et al.*, 1999). Although seed area was significantly smaller in our shoot heated samples when compared to control, there were no significant differences in estimated seed weight (two-way ANOVA on the effects of root and shoot heating, p-values > 0.05, Figure 1d). The decrease in seed area without changes to seed weight with heat treatment is similar to previous reports. For example, in quinoa accession Regalona, night time heat treatment (22°C) resulted in yield losses without changes in average seed weight (Lesjak and Calderini, 2017). Similar to our study, Hinojosa et al. 2018 also found that seed weight was not significantly affected by heat in accession QQ74 (Hinojosa, Matanguihan, *et al.*, 2018).

The number of seeds produced per plant was estimated by dividing total yield (g) by individual seed weight (please see “Main panicle, seed, and whole plant imaging” section in Experimental Procedures for details). There was no significant difference in the estimated individual seed weight with heat treatment, but there was a significant decrease in total seed yield which suggests a change in seed number. Accordingly, there was a significant effect of shoot heating on seed number (two-way ANOVA on the effect of shoot heating, p-value < 0.0001), but no significant effect from root heating on seed number (two-way ANOVA on the effect of root heating, p-value = 0.5933). The estimated seed number was an average of ~79% lower in HS treatment than in control (Wilcoxon rank sum test, p-value = 0.00016) and an average of ~78% lower in HRS treatment than in control (Wilcoxon rank sum test, p-value = 0.00016, Figure 1e). Overall, there were fewer seeds produced per plant and less seed area but no difference in seed weight between plants with heated shoots and control plants. This indicates that the observed yield losses in plants with heated shoots are mainly the result of fewer seeds produced.

### Main and secondary panicles from plants with heated shoots had less yield than those from control plants. Tertiary panicles from HRS plants had higher yield than control tertiary panicles

Quinoa yield is the total seed production from many panicles on each plant. Therefore, we further examined where yield losses were occurring on each plant (i.e. primary, secondary, or tertiary panicles; Figure 2a). The 11-day heat treatment was started at first anthesis of the main panicle, but the panicles produced by a quinoa plant emerge and develop at different times. Therefore, the heat treatment was applied to panicles at different developmental stages and for different durations. To assess the contributions of different panicle types to yield, quinoa panicles were classified into three groups (Figure 2a): 1) the main panicle: the first panicle to emerge, at the top of the plant; 2) secondary panicles: panicles at the tip of each branch, emerging after the main panicle; and 3) tertiary panicles: panicles originating from nodes within branches, emerging after the secondary panicle in the same branch.

**Figure 2.**
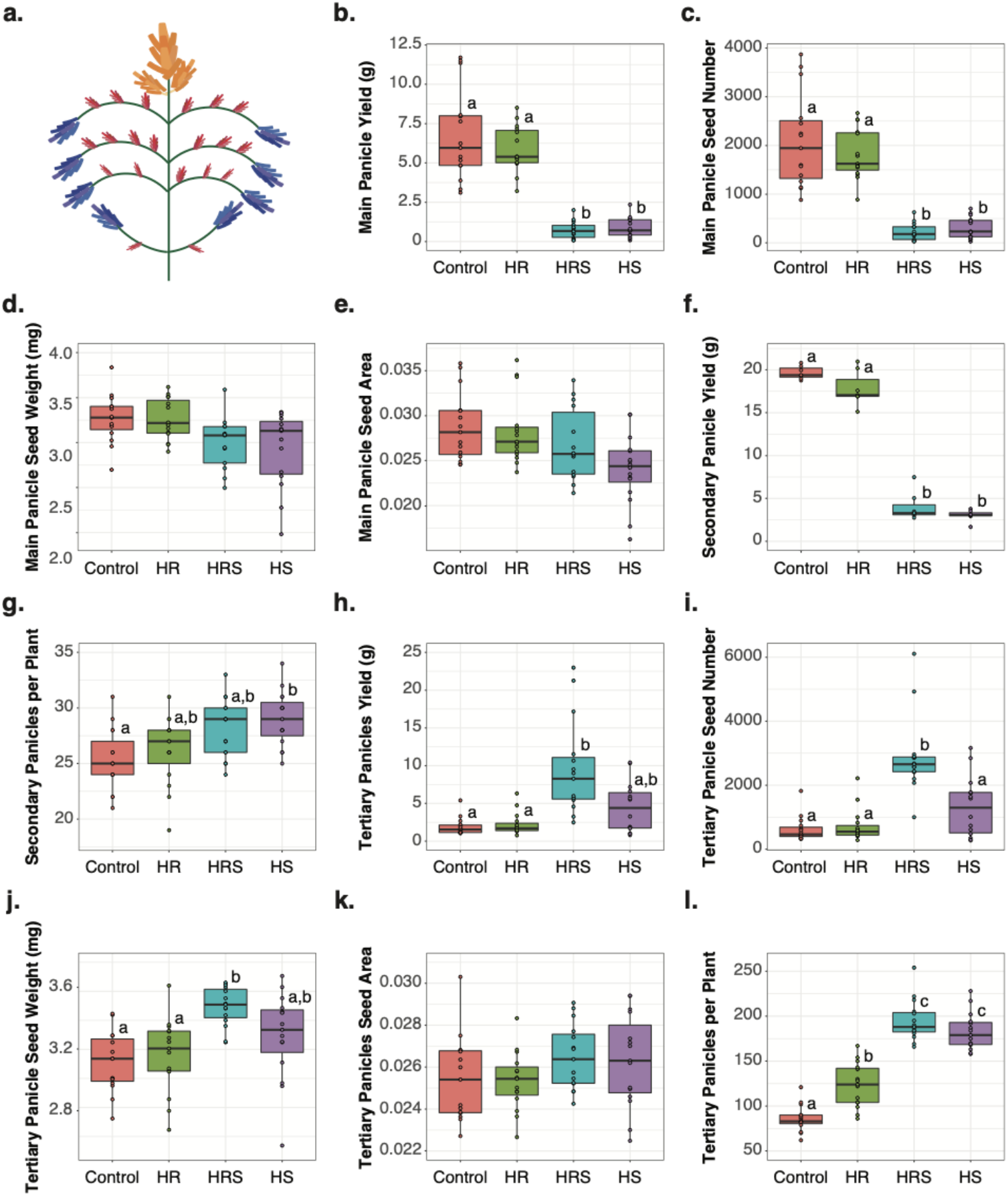
Yield analysis by panicle type. a) Locations of the main panicle (the first panicle to emerge; orange), secondary panicles at the tip of each branch (emerging after the main panicle; blue), and tertiary panicles from nodes within branches (emerging after the secondary panicles; red). b) Yield from the main panicle of each plant for each treatment (n=15). c) Number of seeds produced per main panicle for each treatment (n =15). d) Average seed weight per main panicle for each treatment (n=15). e) Average main panicle normalized seed area per plant for each treatment (n=15). f) Aggregated yield from all secondary panicles in each plant, for each treatment (n=15). g) Number of secondary panicles produced by each plant for each treatment (n=15). h) Total tertiary panicle yield per plant for each treatment (n=15). i) Number of seeds produced from all tertiary panicles in each plant for each treatment (n=15). j) Average tertiary panicle seed weight per plant for each treatment (n=15). k) Average normalized tertiary panicle seed area from each plant and for each treatment (n=15). l) Number of tertiary panicles produced by each plant for each treatment (n=15). Letters above boxes represent statistical significance at p-value < 0.05 from a Wilcoxon rank sum test.

Main panicle yield was significantly affected by shoot heating (two-way ANOVA on the effect of shoot heating, p-value < 0.0001) but not by root heating (two-way ANOVA on the effect of root heating, p-value = 0.6953). Main panicles from HS plants yielded 87% less than control plants (Wilcoxon rank sum test, p-value = 0.00017, Figure 2b). Similarly, main panicles from HRS plants yielded 89% less than control (Wilcoxon rank sum test, p-value = 0.00019, Figure 2b). Based on image analysis, losses in main panicle yield from shoot heating were due to a reduction in the number of seeds produced (Figure 2c). HRS plants produced 89% less seeds than control plants (Wilcoxon rank sum test, p-value < 0.0001) and HS plants produced 85% fewer seeds than control plants (Wilcoxon rank sum test, p-value < 0.0001). Seed number from HR plants was not significantly different from control plants (Wilcoxon rank sum test, p-value < 0.87). Main panicle seed weight (Figure 2d) and area (Figure 2e) were not significantly changed by heat treatments (Wilcoxon rank sum test, p-values > 0.05).

Like main panicle yield, secondary panicle yield was also affected by shoot heating (two-way ANOVA on the effect of shoot heating, p-value < 0.0001) but not by root heating (two-way ANOVA on the effect of root heating, p-value = 0.1801). The average yield of all the secondary panicles from each HS plant was 85% lower than control plants (Wilcoxon rank sum test, p-value = 0.0021, Figure 2f), despite producing 4 more secondary panicles on average than control plants (Wilcoxon rank sum test, p-value = 0.021, Figure 2g). HRS plants also produced an average of 79% less yield from secondary panicles than control plants (Wilcoxon rank sum test, p-value = 0.0021, Figure 2f). Thus, the observed dramatic yield losses from secondary panicles were largely due to shoot heating, similar to main panicles. Unlike HS, HRS plants did not produce significantly more secondary panicles than control plants (Wilcoxon rank sum test, p-value = 0.067).

Unlike main and secondary panicles, tertiary panicles emerged after heat treatment ended. All heat treatments had more tertiary panicles than control. HR plants produced ~43% more tertiary panicles than control plants (Wilcoxon rank sum test, p-value = 0.00026, Figure 2l). HRS plants (Wilcoxon rank sum test, p-value = 0.00002) and HS plants (Wilcoxon rank sum test, p-value = 0.00002) produced more than double the number of tertiary panicles than control plants (Figure 2l). Despite the increase in tertiary panicle number in all heat treatments, increased tertiary panicle yield occurred only in HRS plants but not HR or HS plants. HRS plants tertiary panicles had 5-fold more yield than control tertiary panicles (Wilcoxon rank sum test, p-value = 0.0000057, Figure 2h). The tertiary panicle yield from HS plants (Wilcoxon rank sum test, p-value = 0.18) and HR plants (Wilcoxon rank sum test, p-value = 0.7, Figure 2h) were not significantly different from control plants.

Based on image analysis, the higher yield of tertiary panicles observed in HRS plants was due to higher individual seed weight than control plants (Wilcoxon rank sum test, p-value < 0.0001, Figure 2j), as well as higher number of seeds produced (Wilcoxon rank sum test, p-value < 0.0001, Figure 2i), but not to changes in seed area (Wilcoxon rank sum test, p-values > 0.05, Figure 2k). The increased tertiary panicle yield and seed weight in HRS plants suggests that more resources are allocated to tertiary panicles compared to control. However, this reallocation of resources to tertiary panicles in HRS plants cannot compensate for yield losses in the main and secondary panicles, because HRS plants still produced less total yield than control plants.

### Secondary panicle yield was significantly affected by shoot heating and panicle position in the plant but not by the length of heat treatment

Shoot heating significantly affected the overall yield from all secondary panicles in a quinoa plant. However, secondary panicles from a single quinoa plant emerged and matured at different times, creating differences in their exposure to heat treatment. To assess if yield was affected by different exposure lengths to heat, the yield and time of heat exposure of each secondary panicle was recorded. The distribution of yield among secondary panicles was analyzed through factorial analysis to study the effects of: 1) shoot heating; 2) root heating; 3) secondary panicle position in the plant; and 4) days of heat treatment since secondary panicles showed visible anthers. Factors 3 and 4 were analyzed in separate ANOVAs because they were not independent variables, as secondary panicles closer to the top of the plant will likely mature earlier than panicles closer to the bottom. Main panicles were heat treated from first anthesis, however, anthesis could not be used to measure days of heat exposure in secondary panicles, because anthesis of the secondary panicles was inhibited in plants with heated shoots. Therefore, the days of heat treatment since anthers were visible in the secondary panicle were used to measure the effects heat duration on yield. Secondary panicle yield was affected by shoot heating (three-way ANOVA on the effect of shoot heating, p-value < 0.0001) and position of the secondary panicle (three-way ANOVA on the effect of panicle position, p-value < 0.0001, Figure 3a). Interestingly, the length of heat treatment did not have a significant effect on secondary panicle yield (three-way ANOVA effect of days of heat treatment since visible anthers, p-value = 0.13034; Figure 3b), indicating that a short heat exposure during anthesis may be sufficient to cause significant yield losses.

**Figure 3.**
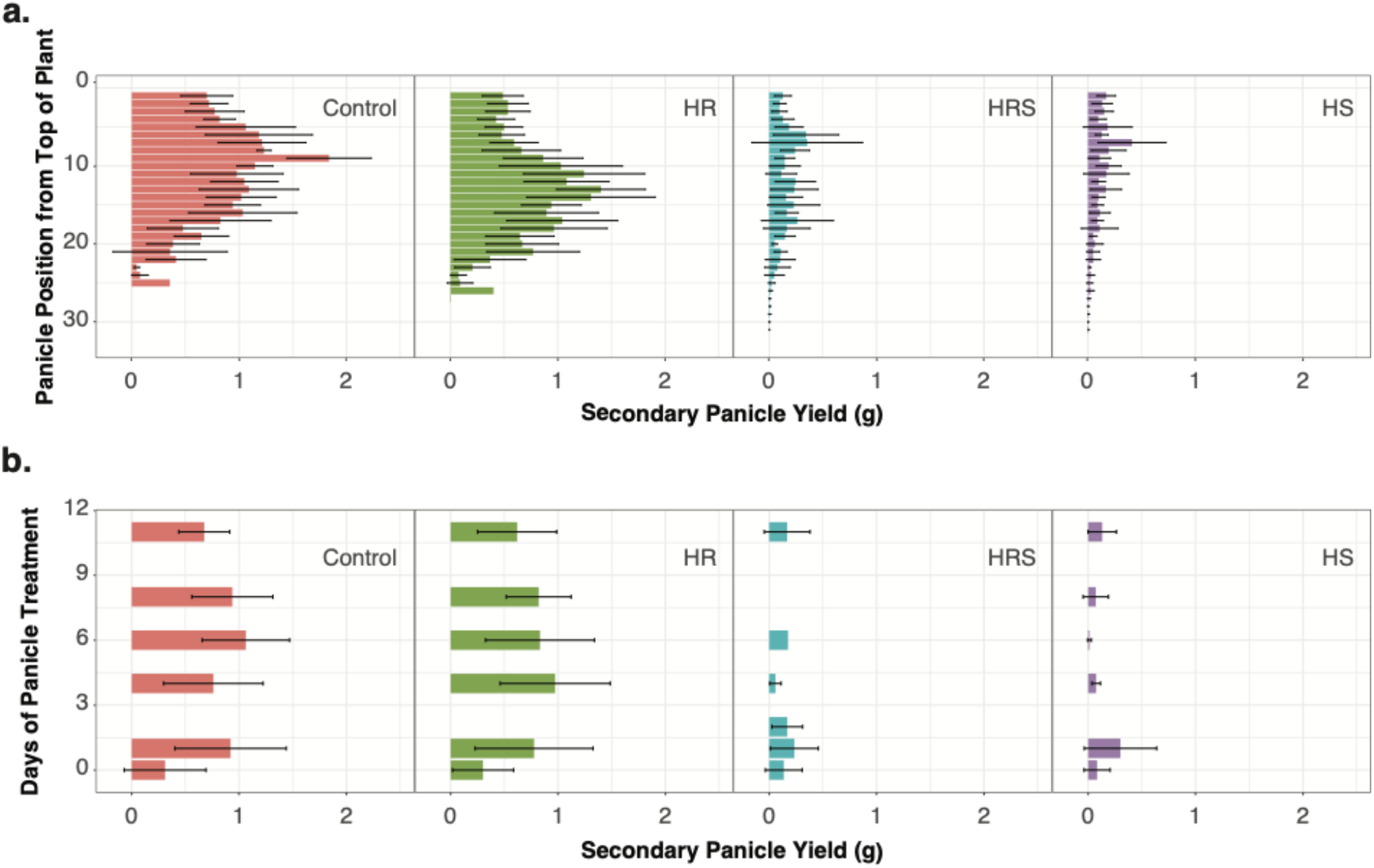
Analysis of the yield of secondary panicles. a) Yield of individual secondary panicles by position in the plant for each treatment (n=1 to 15 secondary panicles, depending on position). Yield from individual secondary panicles heat treated for 0, 1, 4, 6, 8, or 11 days (n=9 panicles per treatment for 0 and for 8 days; n=46 per treatment for 1 day; n=19 per treatment for 4 days; n=59 per treatment for 6 days; n=10 per treatment for 11 days). Error bars represent standard deviation.

### Shoot heating results in significantly fewer flowers developing fruit

To investigate the cause of reduced seed number in HS and HRS plants, flower development in the main panicle was observed during and after heat treatment. After 11 days of heat treatment, all plants were grown at control temperature until harvest. At 2, 24, and 45 days after the heat treatment ended, the total number of flowers and flowers bearing fruit in the apical 5 cm of the main panicle were counted. Fruit production was significantly lower in HS and HRS plants than in control plants (Kolmogorov-Smirnov test, p-values < 0.00001), while root heating did not have a significant effect on fruit production (Kolmogorov-Smirnov test, p-value = 0.09956, Figure 4a). There was a significant effect of shoot heating on fruit production at all time points measured (2, 24, and 45 days after heat treatment ended; two-way ANOVA p-values < 0.00001). HS plants produced ~35.9% of the fruit produced by control plants at 2 days after heat treatment ended (Wilcoxon rank sum test, p-value = 0.02093), ~23.1% at 24 days after heat treatment ended (Wilcoxon rank sum test, p-value < 0.00001), and ~31.4% at 45 days after heat treatment ended (Wilcoxon rank sum test, p-value < 0.00001). HRS plants produced ~16.8% of the fruit produced by control plants at 2 days after heat treatment ended (Wilcoxon rank sum test, p-value = 0.00079), ~26% at 24 days after heat treatment ended (Wilcoxon rank sum test, p-value < 0.00001), and ~30.5% at 45 days after heat ended (Wilcoxon rank sum test, p-value < 0.00001). Overall, fruit production was significantly reduced in treatments with yield losses.

**Figure 4.**
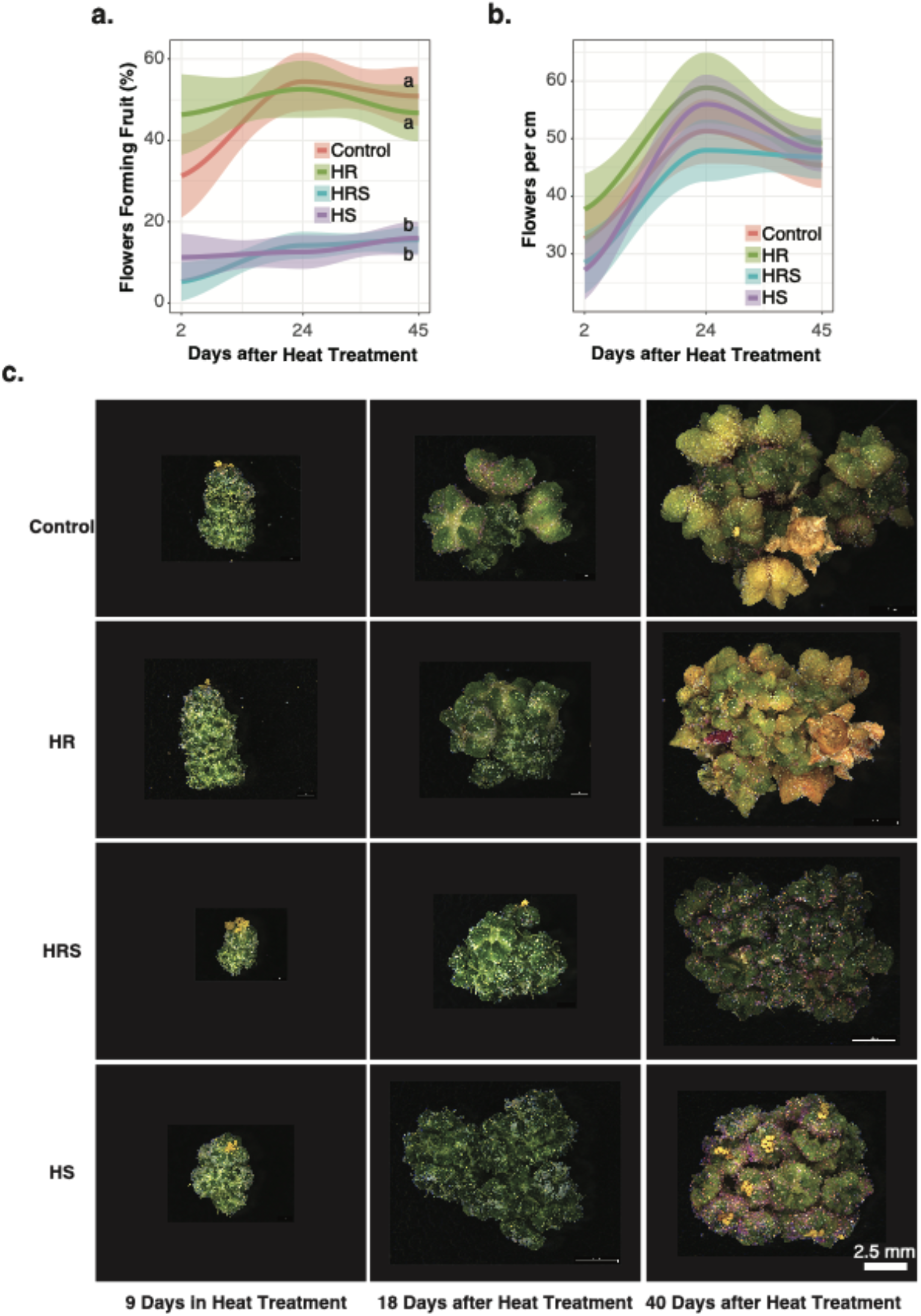
Main panicle flower analysis. a) Fruit production measured on days 2, 24, and 45 after heat treatment ended. b) Flower density in the most apical 5 cm of the main panicle measured on days 2, 24, and 45 after heat treatment ended. Curves resulting from a LOESS polynomial regression are shown (n=30 main panicles, for each timepoint and each treatment). Letters next to curves represent statistical significance at p-value < 0.05 from a Kolmogorov-Smirnov test. c) Flowers from the main panicle for each treatment imaged after 9 days in heat treatment, 18 days after heat treatment ended, and 40 days after heat treatment ended. All images have been scaled to the same scale bar.

Along with fewer flowers producing fruit, another possible contributor to seed yield loss is a lower density of flowers along the panicle. To investigate the possibility that shoot heating lowers the number of flowers produced by a quinoa plant, all the flowers in the apical 5 cm of the main panicle were counted. There were no significant differences in the number of flowers being produced in the apical 5 cm across treatments (Kolmogorov-Smirnov test, p-values > 0.05, Figure 4b). Although, this measurement does not discount the possibility that there are differences in flower concentration in lower regions of the plant, it does suggest that flower concentration in the main panicle is similar between treatments.

To assess possible differences in fruit production and flower development, flowers from the most apical part of the main panicle were observed under the microscope with 9 days of heat treatment, 18 days after heat treatment ended, and 40 days after heat treatment ended. All flowers appeared similar among all treatments with 9 days of heat treatment. However, 18 days after heat treatment ended, several flowers with fruit were visible in both control and HR treatments, but little to no fruit was observed in HS and HRS treatment samples (Figure 4c). At 40 days after heat treatment ended, seed was formed in both control and HR plants, but not in HS or HRS plants (Figure 4c). In general, flowers of HS and HRS plants appeared less developed than control and HR flowers and lacked fruit and seed production (Figure 4c). The lack of fruit production and differences in flower development may explain the observed low seed number and yield loss in treatments with heated shoots.

### Photosynthesis and pollen viability did not show significant changes during or after heat treatment

Since our study observed a significant decrease in yield and fruit production in response to heat stress, we aimed to better understand the physiological changes occurring during heat stress that may have contributed to yield losses. We hypothesized that photosynthesis and pollen viability under heat treatment are negatively impacted. Although changes in photosystem II efficiency have been associated with grain yield (Lin *et al.*, 2018; Ort *et al.*, 2015; Sanchez-Bragado *et al.*, 2016), as well as to responses to heat (Yamamoto, 2016; Mathur and Jajoo, 2014), we did not find evidence that photosystem II efficiency or pollen viability were factors in yield losses after heat treatment (please see supplemental materials for data and details).

### Main panicles had less area and had a more open structure with heated shoot treatment

Next, we hypothesized panicle architecture may be affected by heat. Although image analysis of the main panicles found no significant difference in the height or width of main panicles, main panicle area (two-way ANOVA on the effect of shoot heating, p-value < 0.00001, Figure 5a) and main panicle weight (two-way ANOVA on the effect of shoot heating, p-value < 0.00001, Figure 5b) were reduced after shoot heating relative to control plants. Reduced main panicle area and weight are likely a result of lower fruit production after shoot heating, since fruits are much larger than unfertilized flowers (Figure 5d), and the difference in fruit production from shoot heating was significant. Additionally, solidity, a measure of panicle compactness, was also significantly reduced after shoot heating relative to control plants (two-way ANOVA on the effect of shoot heating, p-value < 0.00001, Figure 5c). Reduced solidity could be caused by main panicles of HS and HRS plants having a more open structure, or by main panicles having fewer branches. To study the cause of reduced solidity, the number of branches in the main panicles were counted but no significant differences were found between any treatment and control (Wilcoxon rank sum test, p-values > 0.05; Figure 5e). This indicates that the observed reduced solidity in plants with heated shoots is likely a result of a more open structure relative to control plants (Figure 5f). Altogether, we found a significant difference in the structure of panicles from heated shoot treatments, which is likely a reflection of differences in fruit production but not flower concentration under heat.

**Figure 5.**
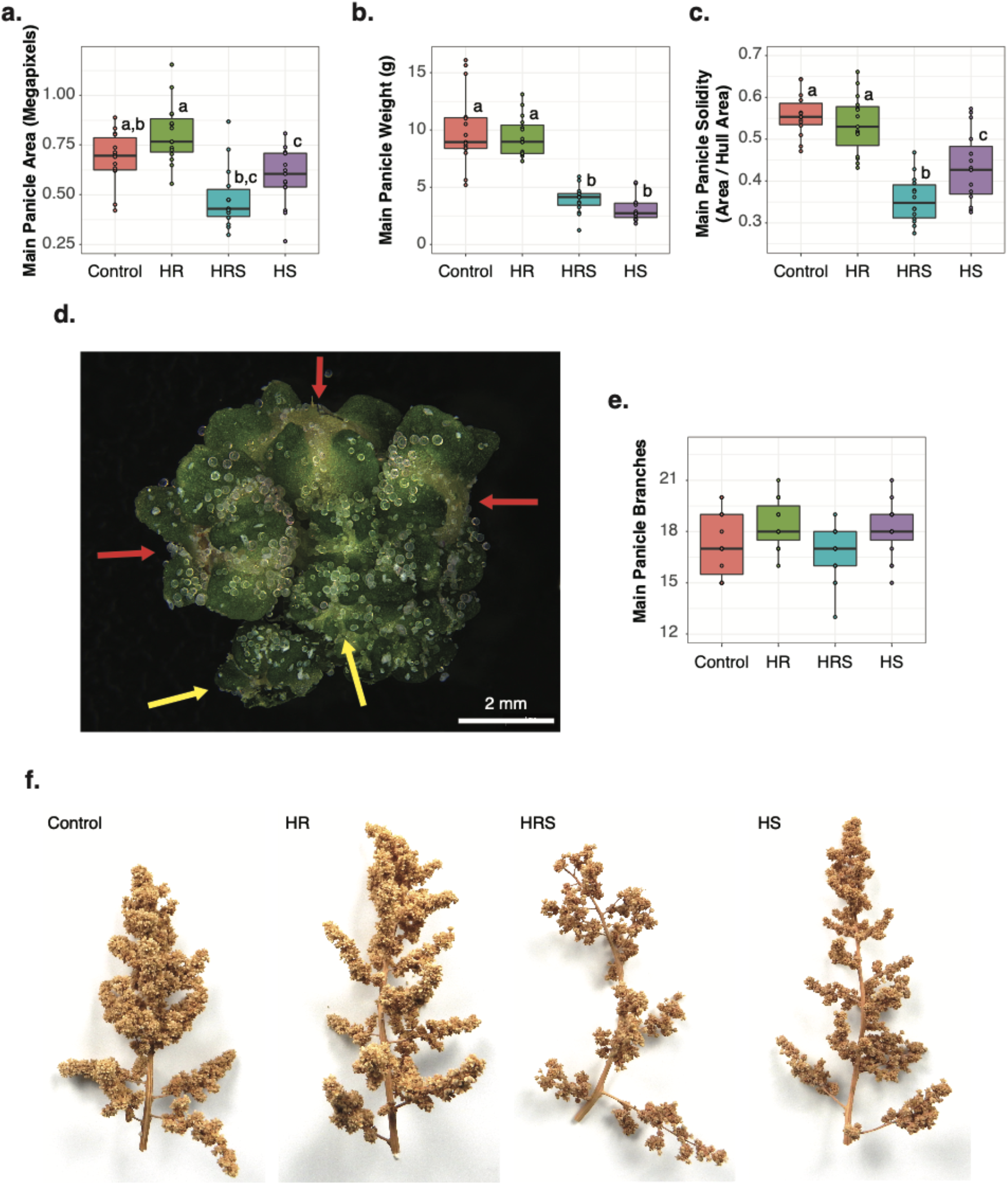
Analysis of main panicle structure. a) Area of each main panicle at harvest per treatment (n=15). b) Weight of each main panicle at harvest for each treatment (n=15). c) Main panicle solidity for each plant and for each treatment (n=15). d) Segment of a quinoa main panicle showing fruit (red arrows) and unfertilized flowers (yellow arrows). e) Number of branches in each main panicle for each treatment (n=15). f) Example images of the main panicle from each treatment group (representative median image for area) at harvest. Letters above boxes represent statistical significance at p-value < 0.05 from a Wilcoxon rank sum test.

### Shoot heating increases the dry weight of non-reproductive shoot biomass

Fruit development was significantly affected in HS and HRS treatments. Therefore, we hypothesized that there were differences in the development of non-reproductive tissues with heat treatment. Quinoa shoot dry weight was measured on the first day of heat treatment, the last day of heat treatment, and at harvest. At the start of heat treatment, plants had similar shoot dry weight (Kruskal-Wallis test, p-value = 0.6227, Figure 6a). After 11 days of heat treatment, shoot dry weight was not significantly different among treatments (Kruskal-Wallis test, p-value = 0.1787, Figure 6a). At harvest, HS plants (Wilcoxon rank sum test, p-value = 0.5929) and HR plants (Wilcoxon rank sum test, p-value = 1) had similar dry weights to control plants (Figure 6a). However, HRS plants had 27% more shoot dry weight than control plants at harvest (Wilcoxon rank sum test, p-value = 0.0042). Interestingly, HRS plants also had significantly more tertiary panicles (Figure 2l) and tertiary panicle yield (Figure 2h), despite having significantly less total yield than control plants (Figures 1a and 1b).

**Figure 6.**
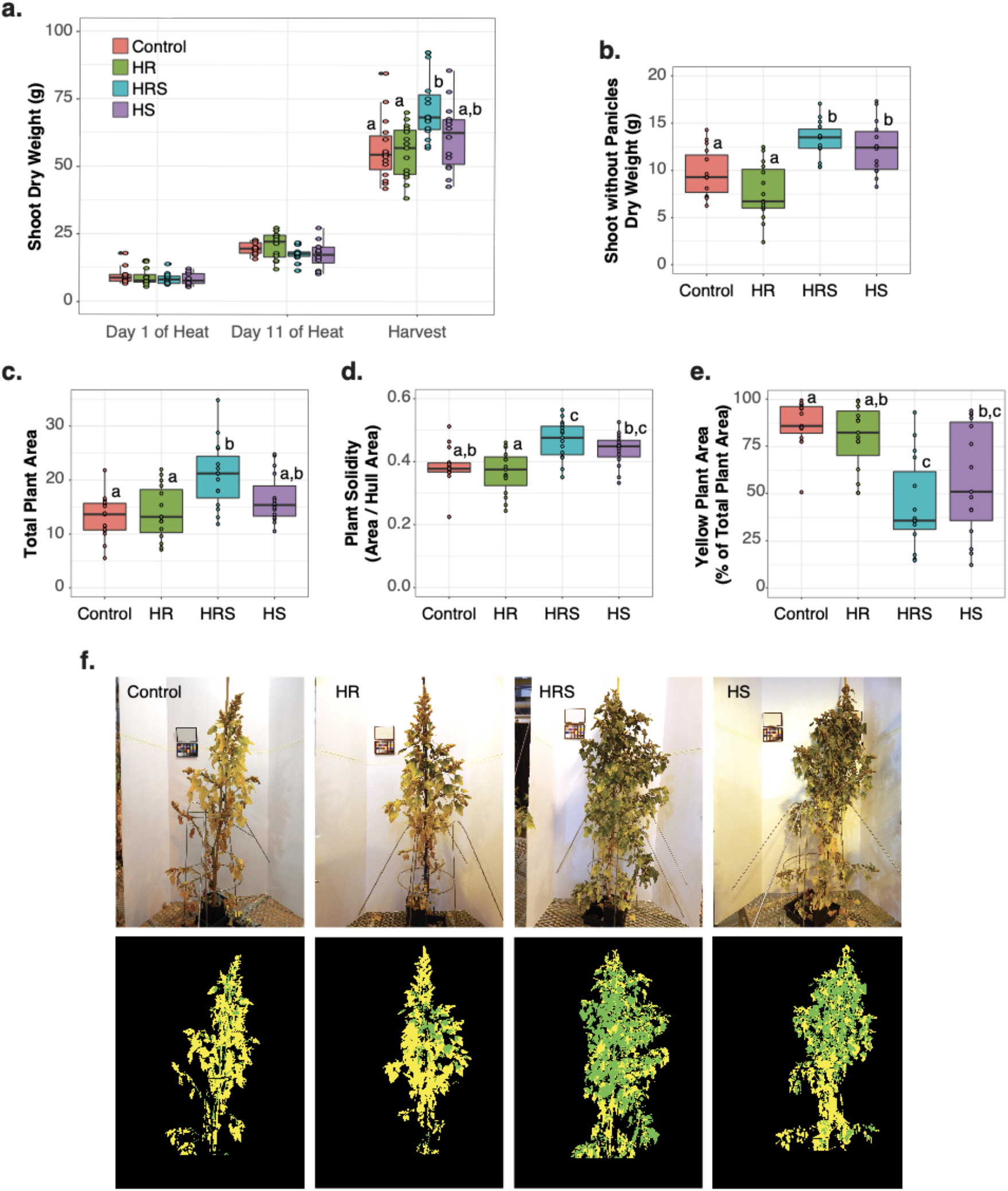
Shoot size and maturity. a) Shoot dry weight (g) with 1 day of heat, 11 days of heat, and at harvest (n=9 for each treatment for days 1 and 11 of heat treatment, and n = 15 for each treatment at harvest). b) Shoot dry weight without panicles at harvest (n=15 for each treatment). Normalized area for each plant and for each treatment (n=15). d) Solidity for each plant and for each treatment (n =15). e) Percent yellow area for each treatment per plant (n=15). f) Example images from each treatment group (representative median image) classified into yellow or green pixels by naive Bayes classifier from PlantCV. Letters above boxes represent statistical significance at p-value < 0.05 from a Wilcoxon rank sum test.

Measurements of total shoot dry weight included the weight of panicles and seeds in addition to leaf and stem tissue. Therefore, to understand if the quantity of non-reproductive biomass was affected by heat, shoot dry weight without panicles was measured. Shoot dry weight without panicles was significantly affected by shoot heating (two-way ANOVA on the effect of shoot heating, p-value < 0.0001) but not by root heating (two-way ANOVA on the effect of root heating, p-value = 0.4997). HS plants had 42% more dry weight in their shoots without panicles than control plants (Wilcoxon rank sum test, p-value = 0.03717), and HRS plants had 48% more dry weight in their shoots without panicles than control plants (Wilcoxon rank sum test, p-value = 0.00242, Figure 6b). Higher shoot dry weight at harvest after heat treatment has also been reported for cultivars Red Head, Cherry Vanilla, Salcedo (Bunce, 2017), and Titicaca (Yang *et al.*, 2016). Interestingly, HS and HRS plants developed more non-reproductive biomass than control plants after the heat treatment concluded, suggesting that heat may trigger irreversible changes in quinoa development.

### Shoot heating delays plant maturity

There was more non-reproductive biomass in HS and HRS plants, therefore we hypothesized that plant maturation was delayed by shoot heat treatment. Analysis of plant images 45 days after heat treatment revealed that plant architecture was affected by shoot heating. In agreement with manually measured shoot dry weight results, plant area extracted from image data was 59.8% higher in HRS plants than in control plants (Wilcoxon rank sum test, p-value = 0.007, Figure 6c). Plants from both HS (Wilcoxon rank sum test, p-value = 0.299) and HR treatments (Wilcoxon rank sum test, p-value = 1) had similar area to control plants (Figure 6c), which mirrors manual shoot dry weight measurements. Plant solidity (compactness) was higher in shoot-heated plants than in control plants (two-way ANOVA on the effect of shoot heating, p-value = 0.0001), while root heating did not significantly change solidity (two-way ANOVA on the effect of root heating, p-value = 0.5822). Quinoa plants shed their leaves at maturity. Therefore, solidity could be a proxy measurement for maturity because solidity likely decreases when leaves are shed. Accordingly, lower solidity could indicate fewer leaves and a more mature plant. HRS plants had significantly higher solidity than control plants (Wilcoxon rank sum test, p-value = 0.0122, Figure 6d), while both HS plants (Wilcoxon rank sum test, p-value = 0.0768) and HR plants (Wilcoxon rank sum test, p-value = 1) did not show a significant change in solidity as compared to control plants (Figure 6d).

We also used color to quantify plant maturity, with “yellow” being more mature and “green” being less mature. Plant area was classified into green and yellow pixel area using the Naive Bayes Classifier in PlantCV (Gehan *et al.*, 2017). Shoot heating had a significant effect on the percent of yellow plant area (two-way ANOVA on the effect of shoot heating, p-value = 0.0001, Figure 6e). Root heating did not have a significant impact on the percent of yellow plant area (two-way ANOVA on the effect of root heating, p-value = 0.3531). HS plants had 57% yellow area and HRS plants had 44% yellow area, while HR plants had 80% yellow area and control plants had 87% yellow area on average (Wilcoxon rank sum test p-values from comparisons between control and: 1. HS 0.0235, 2, HRS 0.001, and HR 0.9965; Figures 6e and 6f). The decreased proportion of yellow plant area in treatments with heated shoots suggests a delay in maturity in comparison to the control plants. Changes in maturity in shoot-heated samples, especially HRS, in combination with increase in tertiary panicle yield and number suggest that this accession of quinoa uses a heat avoidance strategy rather than an escape strategy. In an escape strategy a plant attempts to finish its life cycle (often early flowering) in response to adverse environmental conditions, whereas in an ‘avoidance’ strategy plant development slows down (Shavrukov *et al.*, 2017).

### Differentially expressed genes in treatments with yield losses

To identify candidate genes involved in yield loss caused by heat in quinoa, gene expression was examined. Differentially expressed genes compared to the control treatment were identified by RNA-seq analysis. Leaf samples were collected from all treatments during day 1 and day 11 (last day) of heat treatment. To facilitate the use of the expression data generated in this study, an R Shiny application, “Quinoa Heat Data Explorer,” was developed as a community tool (please see supplemental section for details).

To associate gene expression to yield loss, genes differentially expressed exclusively in the treatments with yield loss (HRS and HS) were further analyzed (Figure 7a). Day 1 samples had 5825 differentially expressed genes in both HRS and HS treatments. Gene ontology (GO) overrepresentation analysis of the 5825 differentially expressed genes found no overrepresented GO terms, as compared to GO term representation in the entire quinoa genome. Day 11 samples had 1001 differentially expressed genes exclusively in both HRS and HS treatments, also with no GO terms overrepresented. Further analysis focused on the subset of 394 genes differentially expressed in both HRS and HS treatments and during both days 1 and 11 of heat treatment (Figure 7b). No GO terms were overrepresented among these 394 genes and Panther GO enrichment analysis did not find any enriched GO terms in any of the gene lists.

**Figure 7.**
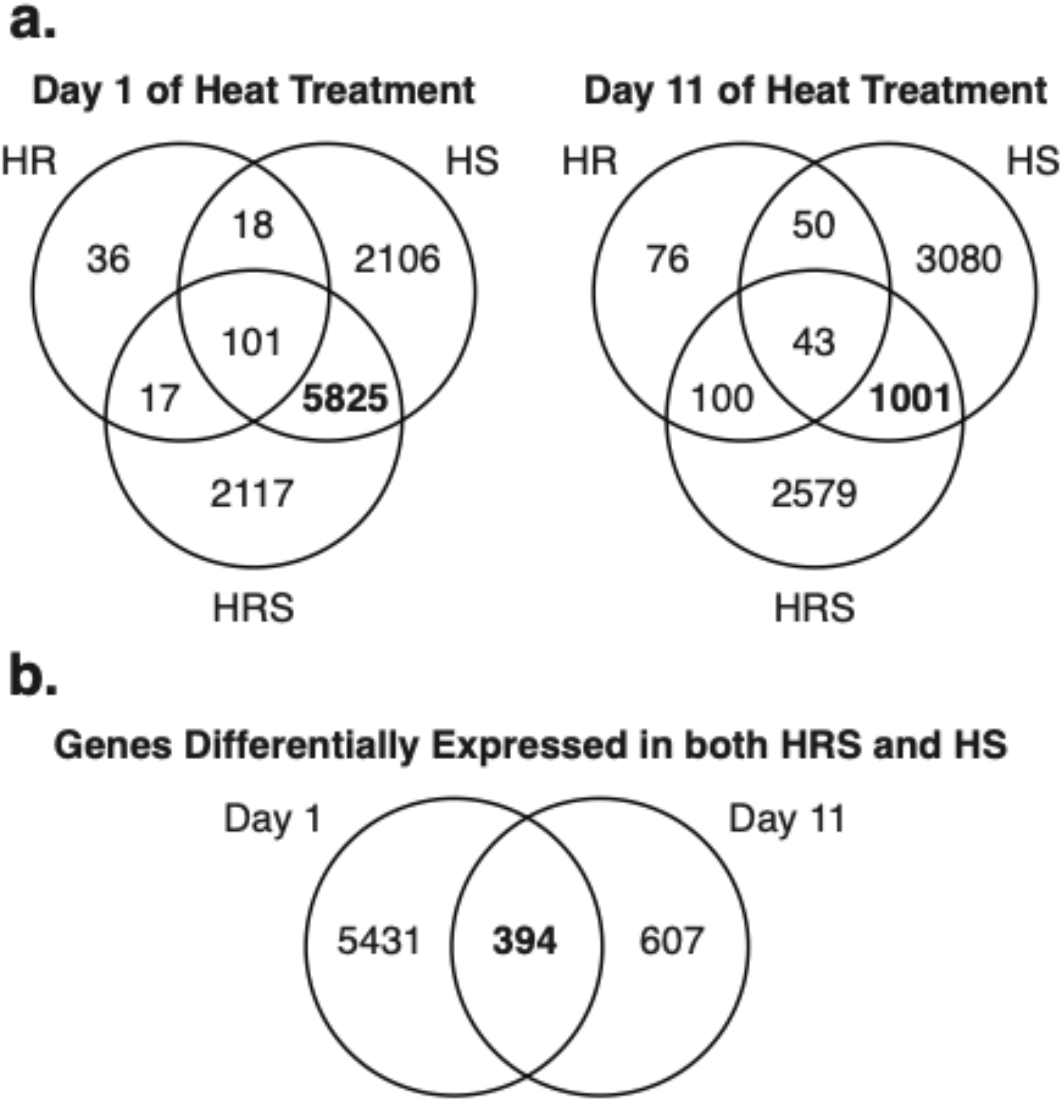
Differentially expressed genes by treatment and day of sampling. a) Venn diagram of differentially expressed genes from control on day 1 of heat treatment and day 11 of heat treatment (n=3 for each timepoint and each treatment). Overlapping genes in treatments with reduced yield are shown in bold. b) Differentially expressed genes overlapping between day 1 and day 11 in treatments with reduced yield (HRS and HS).

### Transcription factor homologs show potential role of flower opening and flower development on yield

A single TF has the potential to modify the expression of many other genes, therefore differentially expressed transcription factors were identified in the 394 genes differentially expressed in HRS and HS treatments. Out of the 394 genes differentially expressed in both HRS and HS treatments during both days 1 and 11 of heat treatment, ten genes were identified as homologous to *A. thaliana* transcription factors. Interestingly, quinoa gene AUR62034763 is a homolog of *AUXIN RESPONSE FACTOR 2* (*ARF2*), a pleiotropic developmental regulator (Okushima *et al.*, 2005) associated with flower opening, fertility, and seed yield in Arabidopsis (Hughes *et al.*, 2008). In Arabidopsis, the *arf2* mutant makes sepals grow longer, preventing petals from separating, so flowers do not open, which lowers seed set and yield (Hughes *et al.*, 2008). Because quinoa flowers have sepals and petals fused (Abdelbar, 2018), AUR62034763 is not likely to function in the same manner as *ARF2* in Arabidopsis. However, identification of this gene led us to examine flower opening and timing during heat treatment. Interestingly, flowers remained closed during the heat treatments with yield losses (HRS and HS) but were open during the day in the treatments with normal yield (HR and control; Figure 8a). A closed flower is likely better protected from heat but less likely to be fertilized. Quinoa inflorescences are composed of hermaphrodite and female flowers, where female flowers depend on the pollen from the hermaphrodite flowers to be fertilized, produce fruit, and ultimately seed (Bhargava *et al.*, 2007b; Abdelbar, 2018). In open flowers, anthers are more spread than in closed flowers (Figure 8b). Flower opening may facilitate pollen dispersal, affecting how many flowers receive pollen and are fertilized. Thus, flowers failing to open could impact fruit formation and therefore, seed number and yield in quinoa under heat.

**Figure 8.**
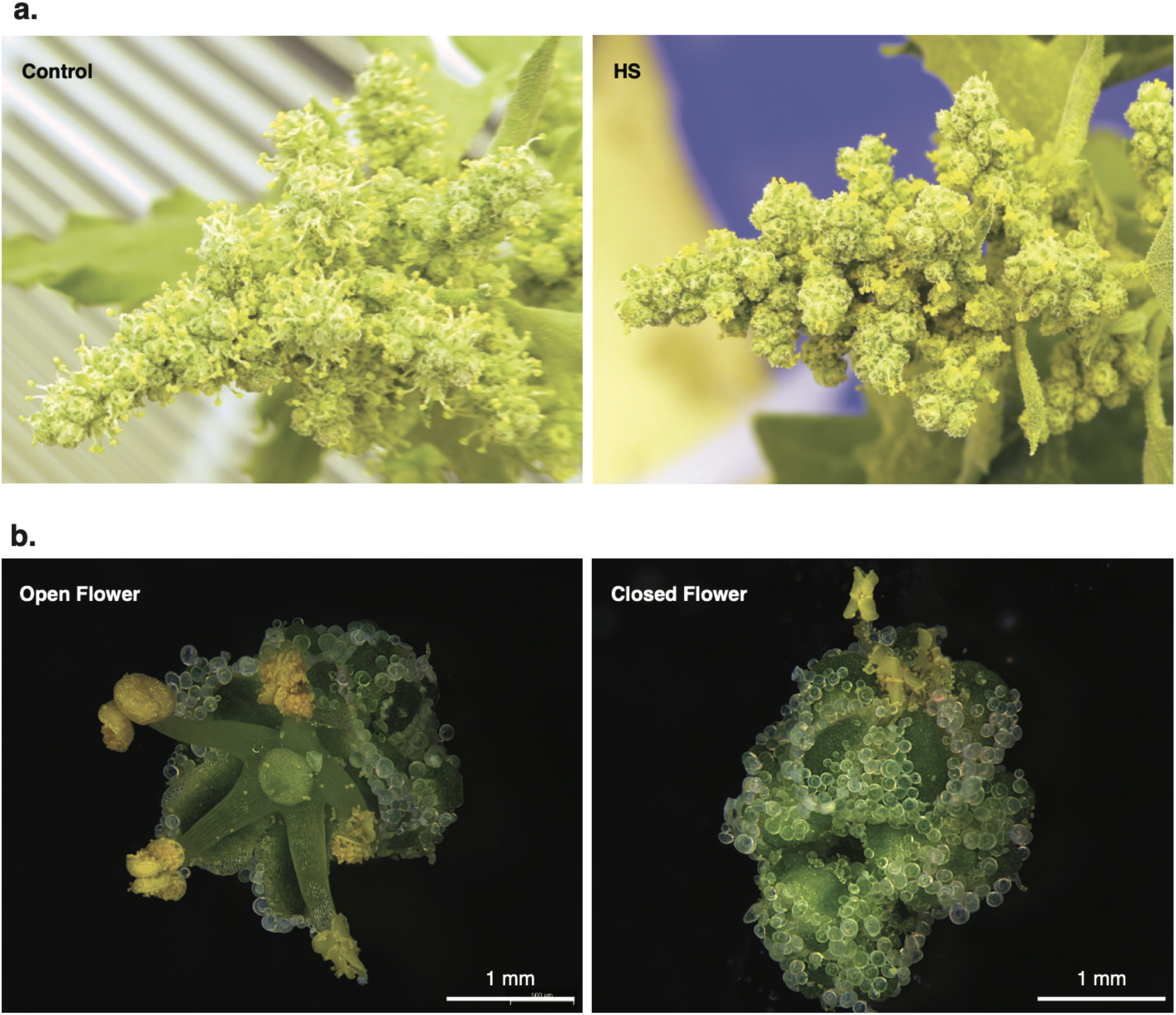
Shoot heating affected floral opening compared to control. a) Representative images of control and heated shoots main panicles. b) Representative images of open and closed quinoa flowers.

Two additional quinoa genes are homologs of TFs associated with flower morphogenesis, and development. AUR62019043 and AUR62033383 are homologs of *AGAMOUS-LIKE 24* (AGL24) and *AGAMOUS-LIKE 14* (AGL14), respectively, which affect flowering and flower development in Arabidopsis (Yu *et al.*, 2004; Torti and Fornara, 2012; Liu *et al.*, 2008; Liu *et al.*, 2007; Thouet *et al.*, 2012; Fernandez *et al.*, 2014; Agliassa *et al.*, 2018; Pérez-Ruiz *et al.*, 2015). Further, *AGAMOUS-LIKE 24* (AGL24) has been shown to be temperature responsive in Arabidopsis (Gregis *et al.*, 2006). AUR62019043 and AUR62033383 have the potential to affect flower development, therefore they could be affecting fruit production, and ultimately yield under heat in quinoa. AUR62008425 is a homolog of *cdf3*, an Arabidopsis TF that affects flowering time under abiotic stress (Corrales *et al.*, 2017). In total, four of the ten TFs differentially expressed in both treatments with yield losses during both days 1 and 11 of heat treatment have been associated with flowering and flower development in Arabidopsis. Since flower development was delayed in quinoa treatments with yield losses (HS and HRS; Figure 4c) and four transcription factors differentially expressed in these treatments are associated with flower development, it is possible that flower development may have a significant role in yield losses with shoot heat treatments. Further, it is possible that one or more of these genes may be involved in these yield losses.

## Conclusions

We explored the physiological changes that occur in quinoa accession QQ74 during and after heat treatment. The shoot heat treatments had the most physiological changes in response to heat. In particular, we find significant reductions in fruit production, delays in maturity and increases in tertiary panicle yield and number in shoot treatments, suggesting that a heat treatment during anthesis triggers an avoidance strategy, prioritizing growth over development, with more resources allocated to tertiary panicles. The increase in tertiary panicle yield at the expense of primary and secondary panicle yield in heated shoot plants compared to control plants, could be an effective survival strategy, but from an agricultural standpoint yield from tertiary panicles would likely not have a significant contribution to total yield because quinoa is typically harvested at main panicle maturity. That being said, as breeding programs commonly select against the development of tertiary and secondary panicles so that more resources are dedicated to the main panicle, they may be inadvertently reducing heat stress tolerance mechanisms in quinoa. In the future, we are still very interested to test if there are differences in seed quality among the treatments (e.g. amino acid profiles) and if there are differences in quality from the different panicles types.

Gene expression data led us to examine floral phenotypes more closely, and we also observed a change in flower opening in the main panicle of HRS and HS treated samples; with flowers remaining closed in the afternoon when control and HR samples had open flowers. This observation fits with the decreased fruit production in the main panicles of HRS and HS samples. Altogether, this work identifies key phenotypes to potentially mitigate heat induced yield decreases in quinoa. For example, if yield decreases of the main panicle are due to floral closures induced by heat then identifying natural variation or screening mutant populations for changes to these phenotypes might be targeted.

## Experimental procedures

### Plant material and growth conditions

QQ74 (PI 614886) seeds were planted in Pro-Mix FPX (Premier Tech Horticulture, Quebec, Canada) soil and grown at 22 °C, 12 hour day photoperiod, 50% relative humidity, 400 µmol m^−2^ s^−1^ light intensity. Plantlets at the 2 to 4 leaves developmental stage were transplanted to Berger BM7 soil (Berger, Quebec, Canada) in 4.5-inch pots, and grown under the same conditions. Before flowering, 60 plants most similar in height and developmental stage were selected, and randomly divided into four treatment groups. Potted plants were placed in 245 cm length × 68 cm width × 16.5 cm height wooden boxes filled with Turface Athletics MVP coarse sand (Profile Products LLC, Buffalo Grove, Illinois) to buffer soil temperature. The soil was covered with a blue mesh (Con-Tact Brand, Pomona, California, USA) to facilitate image processing. When more than 50% of the plants had open flowers (35 to 40-days after planting), all plants were moved to a secondary growth chamber at the same conditions for 2 days, while temperature was adjusted and stabilized for heat treatments. More information about the treatment group boxes is below in the “Heat Treatments” section.

### Heat treatments

Four treatment groups were used in this experiment: 1) Control, with plants growing at 22 °C; 2) Heated roots, with roots growing at 30 °C and shoots growing at 22 °C; 3) Heated shoots, with roots growing at 22 °C and shoots growing at 35 °C; and 4) Heated roots and shoots, with roots growing at 30 °C and shoots growing at 35 °C (Figure S1). To set up for the four treatments, the wooden boxes were split into 2 growth chambers. The first growth chamber was set to control conditions of 22 °C, 12 hour day photoperiod, 50% relative humidity, 400 µmol m^−2^ s^−1^ light intensity, while a second growth chamber was set to 35 °C, 12 hour day photoperiod, 50% relative humidity, 400 µmol m^−2^ s^−1^ light intensity. In the first growth chamber, a heating coil at 30°C was passed through the sand of one wooden box at 2 heights, approximately equally spaced around the pots, to heat the soil to 30 °C (HR). In the second growth chamber at higher temperature, a cool water line running 15.5°C water was passed through the sand of one wooden box at 2 heights, approximately equally spaced around the pots to cool the soil to 22 °C (HS, Figure S1). At zeitgeber time (ZT) 0, plants were divided into four treatments, where each treatment was contained in a wooden box. Heat treatment lasted 11 days. After 11 days of treatment, plants were moved for a day to a third growth chamber at control conditions while temperature was adjusted back to control conditions in the treatment chambers. Once control conditions (22 °C) were reached in treatment chambers, plants were replaced in their respective wooden boxes at control conditions, for 7 to 10 more days, and subsequently moved to a greenhouse at control conditions, until harvest for yield measurements.

### Sampling for pollen viability, fresh weight, and dry weight

Samples from 2 to 4 plants per treatment were collected between ZT2 and ZT4, at days 1 and 11 of heat treatment. Flowers with visible anthers were cut and placed in Alexander stain (Alexander, 1969) for pollen viability assays. Sampled plants were cut to separate shoots from roots. Shoots were immediately weighed to obtain fresh weight, and subsequently both roots and shoots were dried at 40 °C, 30% relative humidity for 3 days and weighed to obtain dry weight.

### Seed harvesting

When plants stopped uptaking water from the soil, watering was stopped and plants were allowed to dry until ready to harvest. Shoots and panicles were cut and stored in paper bags at room temperature (approximately 23°C). Seed was harvested from the shoots using an air blast seed cleaner (ABSC; Almaco, Nevada, Iowa). Seed was then manually cleaned with a mesh and stored in paper envelopes at room temperature.

### Main panicle, seed and whole plant imaging

A Raspberry Pi computer controlling a SLR camera (Nikon COOLPIX L830) on a camera stand was used to collect main panicle and seed images (https://doi.org/10.5281/zenodo.3352281). The camera setup to collect seed images is described in detail in Appendix 3 of Tovar et al. 2018 (Tovar *et al.*, 2018). Main panicle images were analyzed using PlantCV (Gehan *et al.*, 2017) to measure main panicle area, width, height, hull area, solidity, perimeter, longest axis, and hull vertices. The python script analyze_image.py (https://github.com/danforthcenter/quinoa-heat-tovar/blob/master/analyze_image.py) was run over all images using the PlantCV script pantcv-pipeline.py. The R script used to analyze the main panicle output measurements is available on Github (https://github.com/danforthcenter/quinoa-heat-tovar/blob/master/main_panicle_image_data_analysis.R). To estimate seed size, a subset of quinoa seeds collected from control, heated roots, heated shoots, and heated roots and shoots treated plants were imaged and then that subset of seeds was weighed. The number of seeds produced per plant was estimated by dividing the total seed yield of every plant by the estimated individual seed weight of the same plant. A 0.5 inch tough spot was used as a size marker for seed images. Images were analyzed using PlantCV (Gehan *et al.*, 2017) to measure seed area. Available on Github are an example Jupyter notebook for seed image processing (https://github.com/danforthcenter/quinoa-heat-tovar/blob/master/seed-image-analysis/plantcv-seed-phenotyping-phenomatics.ipynb), and the R script used to process seed area results (https://github.com/danforthcenter/quinoa-heat-tovar/blob/master/seed-image-analysis/heat-seed-quinoa-analysis.R). Adjustments to the region of interest were made for different seed images, to make sure that all seed and size marker were included in the analysis.

Whole plant images were acquired using the same SLR camera used for main panicle and seed imaging, with white poster boards as background (https://doi.org/10.5281/zenodo.3352281). Whole plant images were acquired 45 days after the end of heat treatment (approximately 95 days after planting). An example Jupyter notebook used to analyze whole plant images (https://github.com/danforthcenter/quinoa-heat-tovar/blob/master/whole_plant_images.R) and the probability density functions file (https://github.com/danforthcenter/quinoa-heat-tovar/blob/master/whole_plant_pdfs.txt) used to classify green and yellow plant parts are available on Github. Adjustments to the placement of black boxes, thresholds, and the region of interest were made in the Jupyter notebook when analyzing whole plant images, to make sure that the entire plant was included in the analysis. The side of the pot was measured in pixels using ImageJ (Schneider *et al.*, 2012) for each whole plant image, and used to normalize linear measurements (perimeter, height, width, and longest axis), while the square of the pot side was used to normalize area measurements (area, and hull area). The R script used to analyze whole plant image measurements is available on Github (https://github.com/danforthcenter/quinoa-heat-tovar/blob/master/whole_plant_images.R).

### RNA-Seq

Samples were collected from 3 plants per treatment at four time points: 1, 2, and 11 days of heat treatment, and 1 day after heat treatment ended. Each plant was considered a biological replicate. Leaf tissue, root tissue, and flower tissue was collected at zeitgeber time 2 and immediately frozen in liquid nitrogen. Although four time points and multiple tissue types were collected, only leaf samples from day 1 and day 11 were processed further for RNA-seq. Total RNA was extracted from quinoa leaves with the RNeasy Plant Mini Kit, including the RNase-free DNase Set, as recommended by the manufacturer (Qiagen, Hilden, Germany). RNA was quantified with Qubit RNA BR Assay Kit (Invitrogen, Carlsbad, California), and quality was assessed with the Agilent 6000 Nano Kit (Agilent, Santa Clara, California). Strand-specific library construction and paired-end RNA sequencing were done by Novogene (Chula Vista, California) using Illumina HiSeq 4000 (San Diego, California). Resulting paired-end reads were analyzed for quality using FastQC version 0.11.7 (Andrews *et al.*, 2012), where the “Per base sequence content” showed the first 10 to 20 bases on the 5’ ends contained low quality reads, suggesting the need to trim those bases. Based on FastQC quality analysis, reads were trimmed using Trimmomatic version 0.38 (Bolger *et al.*, 2014) with Phred 33 quality scores (option “-phred33”), removing adapters (option “ILLUMINACLIP:TruSeq3-PE.fa:2:30:10”), cropping 15 bases from the 5’ end (option “HEADCROP:15”), and using the default options “LEADING:3 TRAILING:3 SLIDINGWINDOW:4:15 MINLEN:36”. After read trimming, the “Per base sequence content” in FastQC showed good quality, indicating reads were ready for gene expression profiling. A quinoa transcriptome index was created with Kallisto (Bray *et al.*, 2016) version 0.44.0, using the “kallisto index” command with required argument “-i” and no optional arguments, from the published quinoa transcriptome (file Cquinoa_392_v1.0.transcript.fa.gz) (Jarvis *et al.*, 2017) available at Phytozome (Goodstein *et al.*, 2012). Transcript abundance was quantified with Kallisto version 0.44.0 (Bray *et al.*, 2016) by pseudo aligning the trimmed paired-end reads to the created quinoa transcriptome index, using the “kallisto quant” command with required arguments “-i” and “-o”, and optional bootstrapping set to 100 (argument “-b”). Differential analysis of transcript levels between treatments and control was done with Sleuth version 0.30.0 (Pimentel *et al.*, 2017) in RStudio version 1.1.463 and R version 3.5.2. A q-value < 0.05 was used to call differentially expressed genes. The raw RNA-seq reads and the relative gene expression levels resulting from differential expression analysis with Sleuth are available at NCBI GEO submission GSE128155. The R script rna_seq_quinoa_heat.R used to analyze differential gene expression is available at https://github.com/danforthcenter/quinoa-heat-tovar/blob/master/rna_seq_quinoa_heat.R.

### Gene ontology analysis

Gene ontology (GO) was performed on differentially expressed genes found in RNA-seq analysis. Panther 14.0 (http://pantherdb.org/) was used for GO analysis (Mi *et al.*, 2013). Differentially expressed gene lists were associated to their Panther IDs from the quinoa genome (v1.0) annotation (Jarvis *et al.*, 2017), available at Phytozome (Goodstein *et al.*, 2012). Text files with gene lists were uploaded into Panther using the “PANTHER Generic Mapping” option, following Panther recommended file formatting, with quinoa gene names on the first column, and Panther IDs on the second column, and using “NOHIT” whenever a gene did not have an associated Panther ID. The b values obtained from differential expression analysis with Sleuth version 0.30.0 were used as the third column (numerical value of the experiment) for statistical enrichment tests. Statistical overrepresentation and enrichment tests were performed using Fisher’s Exact test and false discovery rate (FDR) p-value correction, on all 3 GO categories: molecular function, biological process, and cellular component. For statistical overrepresentation tests, a text file containing all genes in the quinoa genome as the first column and their respective Panther IDs in the second column was uploaded and used as a Reference List. The R script GO_analysis_quinoa_heat.R used to produce the files used for GO analysis is available at https://github.com/danforthcenter/quinoa-heat-tovar/blob/master/GO_analysis_quinoa_heat.R.

### Transcription factor homolog identification

Differentially expressed genes obtained from RNA-seq analysis were screened to identify homologs of *Arabidopsis thaliana* transcription factors. *A. thaliana* orthologs corresponding to each quinoa gene were obtained from the quinoa genome annotation (Jarvis *et al.*, 2017) available in Phytozome (Goodstein *et al.*, 2012). To identify quinoa transcription factor homologs, the list of quinoa genes and their respective *A. thaliana* gene homolog names were cross referenced with the list of Arabidopsis transcription factors available at http://planttfdb.cbi.pku.edu.cn/ (Jin *et al.*, 2017). The R script used to identify transcription factor homologs is included in the rna_seq_quinoa_heat.R R script, available at https://github.com/danforthcenter/quinoa-heat-tovar/blob/master/rna_seq_quinoa_heat.R.

### Statistical analysis

Data was analyzed in RStudio version 1.1.463, with R version 3.5.2. A p-value < 0.05 was considered statistically significant. Each plant was considered a biological replicate. Curves were analyzed using a Kolmogorov-Smirnov test (Kolmogorov, 1933; Smirnov, 1948) with the ks.test function (Figures 4a, 4b, S2a, and S3). Individual timepoints were analyzed with a robust 2-way ANOVA using the med2way function from the WRS2 package (https://r-forge.r-project.org/projects/psychor/), with factors: 1) Root heating and 2) Shoot heating. When a statistical effect was found in the ANOVA, a Kruskal-Wallis test (Kruskal and Wallis, 1952) was performed with the kruskal.test function, to confirm differences existed among treatments. If differences were confirmed by the Kruskal-Wallis test, a pairwise Wilcoxon rank sum test (Mann and Whitney, 1947) using the pairwise.wilcox.test function with a Benjamini and Yekutieli p-value correction (Benjamini and Yekutieli, 2001) (argument p.adjust.method = "BY") was used to find which treatment was different from which, and the respective p-values (Figures 1, 2, 5, 6, and S2b). Analysis of individual secondary panicle yield was done through ANOVAs with factors: 1) Shoot heating, 2) Root heating, and 3) Panicle position (Figure 3a), or 1) Shoot heating, 2) Root heating, and 3) Length of heat treatment (Figure 3b). For RNA-seq data analysis (Figure 7), please see the *RNA-seq* section of Experimental procedures.

### Data Statement

All data from this study is publicly available. The public locations for each data set are mentioned in the Experimental procedures section.

## Acknowledgements

We would like to thank Rick Jellen and Jeff Maughan (BYU) for seed resources. Thank you also to Elizabeth Kellog and Yunqing Yu for guidance on inflorescence and flower anatomy and development, as well as on flower microscopy imaging. Keith Slotkin provided helpful suggestions for editing our manuscript. Teng Chong provided guidance on pollen viability and Alexander staining assays. Zoee Perrine helped provide context to photosynthesis results. Mark Wilson gave bioinformatics suggestions that were helpful to complete differential gene expression analysis. Jeffrey Berry gave suggestions on statistical data analysis. Kristina Haines, Stephen Kraeuter, Kevin Reilly, and William Kezele provided growth chamber and greenhouse plant care and support. This research was supported by the Donald Danforth Plant Science Center.

## Supporting Methods

### Photosynthetic measurements

Photosynthetic rates were measured with a MultispeQ (version 1.0, PhotosynQ, East Lansing, Michigan), using the Leaf Photosynthesis MultispeQ V1.0 (http://photosynq.org/protocols/leaf-photosynthesis-multispeq-v1-0, (Kuhlgert *et al.*, 2016)) and The One v3.0 (http://photosynq.org/protocols/the-one-v3-0-phi2-npqt-using-multi-phase-flash-with-qi-qe-phi2no-rg) protocols. MultispeQ measurements were taken on 3 plants per treatment, every day during the 11 days of heat treatment, between ZT8 and ZT10. Leaves of the same developmental age were measured every day. Fv/Fm was measured on day 11 of heat exposure, using a Li-cor LI-6400XT (Li-cor, Nebraska, USA) following manufacturer’s recommendations.

### Pollen viability

At least two anthers from different flowers of the same plant were manually opened to release pollen into 10 µl of Alexander stain (Alexander, 1969). Pollen was stained for 15 minutes and then centrifuged at 20,000 × g for 1 minute to spin pollen down. To concentrate pollen in the stain, the top 5 µl of stain were discarded. Pollen was resuspended in the remaining 5 µl of stain and analyzed for viability under a brightfield microscope (Omax, Gyeonggi-do, Korea). At least 100 pollen grains were counted per plant to estimate percentage viability, considering green-stained pollen as non-viable, and red or purple-stained pollen as viable (Alexander, 1969).

## Supporting Information

### Photosystem II efficiency is not different before, during, or after heat treatment

To assess the effect of heat on photosystem II efficiency in quinoa, Phi2, a measurement of the quantum yield of photosystem II, was measured using a MultispeQ (v1.0) (Kuhlgert *et al.*, 2016). Measurements were taken from 11 days before heat treatment started, during the 11 days of heat treatment, and 8 days after heat treatment ended. Analysis of Phi2 data using the Kolmogorov-Smirnov test showed that the heated shoots treatment was different from control (p-values 0.003688, Figure S2a). To verify the differences in photosystem II efficiency measured with Phi2, Fv/Fm was used as an independent measurement of the quantum yield of photosystem II. Fv/Fm was measured on day 11 of heat treatment, when we might expect the impact of the progressive heat treatment to be the greatest (Becker *et al.*, 2017; Yang *et al.*, 2016). However, there was no statistically significant effect of heat on Fv/Fm in any treatment (Kruskal-Wallis test p-value 0.09369, Figure S2b). Similarly, Hinojosa et al. 2018 (Hinojosa, Matanguihan, *et al.*, 2018) also found no effects of heat on Fv/Fm in quinoa accession QQ74. A different study in quinoa cultivar Titicaca found that plants grown at 25/20°C (day/night) had higher Fv/Fm than plants grown at 18/8°C (day/night) (Yang *et al.*, 2016), but the yield from these plants was not reported. Overall, there was no conclusive evidence to support changes in photosystem II efficiency due to heat in quinoa. This likely indicates that photosystem II is not a significant factor in yield losses of quinoa from heat.

### Pollen viability did not change after heat treatment

The observed low fruit production after shoot heating could be a result of lower pollen viability under heat treatment, therefore pollen viability was measured. Pollen viability was assessed at the start and end of heat treatments by measuring the rate of pollen abortion through Alexander staining (Alexander, 1969). Pollen viability as measured by pollen abortion in shoot or root heated samples was not significantly affected as compared to control (Kolmogorov-Smirnov test p-values > 0.05, Figure S3). Since heat did not significantly affect the rate of aborted pollen, this would suggest that the observed low fruit production after shoot heating is unlikely due to changes in pollen viability. Interestingly, Hinojosa et al. 2018 (Hinojosa, Matanguihan, *et al.*, 2018) found heat resulted in 63% lower pollen viability than control, in the same quinoa cultivar used in this study (QQ74). However a different pollen staining method (tetrazolium), was used (Hinojosa, Matanguihan, *et al.*, 2018). Interestingly, the significantly lower pollen viability in Hinojosa et al. 2018 did not affect yield (Hinojosa, Matanguihan, *et al.*, 2018). Since no differences in pollen viability were found from heat treatment, and the study of Hinojosa et al. 2018 (Hinojosa, Matanguihan, *et al.*, 2018) found that a dramatically lower pollen viability did not affect yield, this would suggest that pollen viability is not the main limitation for quinoa fruit and seed production. However, the different pollen viability results of this study and of Hinojosa et al. 2018 (Hinojosa, Matanguihan, *et al.*, 2018) highlight the importance of the method used to measure pollen viability. A more definitive method for measuring pollen viability is in vitro pollen germination (Sato *et al.*, 2000; Hinojosa, Matanguihan, *et al.*, 2018), but to our knowledge, there is no published in vitro pollen germination protocol for quinoa (Hinojosa, Matanguihan, *et al.*, 2018). Pollen viability studies through in vitro germination would provide more conclusive evidence to whether pollen viability is affected by heat treatment in quinoa, and its potential impact in fruit and seed production.

### Shoot fresh weight did not show significant changes during heat treatment

To assess the effect of heat on shoot biomass, shoot fresh weight was measured on the first and last days of heat treatment. At the start of heat treatment shoot fresh weight was very similar among all plants (Kruskal-Wallis test p-value 0.9465). On the last day of heat treatment, shoot fresh weight had not significantly changed (Kruskal-Wallis test p-value 0.1226, Figure S4a). To assess differences in water content, the amount of water in each plant was also calculated by subtracting the shoot dry weight from the shoot fresh weight of each plant. As with fresh weight, no differences in water content were found neither on the first (Kruskal-Wallis test p-value 0.8428) nor on the last day of heat (Kruskal-Wallis test p-value 0.1086, Figure S4b). Since it was expected that heat would increase water demands for treated plants, plants were well watered according to demand to prevent drought stress. Thus, quinoa was able to retain control water levels even during heat treatment.

### Root dry weight did not show significant changes during heat treatment

Root dry weight was measured on the first and last days of heat treatment to assess the effects of heat on root biomass. There were no significant differences in root dry weight, both at the start (Kruskal-Wallis test p-value 0.3462) and end of heat treatment (Kruskal-Wallis test p-value 0.3351, Figure S5). The study of Hinojosa et al., 2018 (Hinojosa, Matanguihan, *et al.*, 2018) also found no significant differences in root dry weight 8 days after heat treatment, and therefore also suggests that heat does not affect root biomass in quinoa. Although there was no indication of changes in root dry weight from heat, in both studies, plants were grown in pots, which could have limited root growth. Measurements of root dry weight from heat-treated plants grown in the field would provide a more conclusive assessment of root dry weight changes after heat.

### Quinoa Heat Data Explorer: a tool for navigating and analyzing quinoa differential gene expression under heat

Quinoa Heat Data Explorer is a tool built to facilitate sharing and analysis of the gene expression data obtained in this study with the quinoa research community. Quinoa Heat Data Explorer can be accessed at http://shiny.datasci.danforthcenter.org/quinoa-heat/, or downloaded from https://github.com/danforthcenter/quinoa-heat-tovar and run locally using R or RStudio. With the Quinoa Heat Data Explorer tool, users can browse and search differentially expressed quinoa genes using significance level cutoffs (q-values). Alternatively, users can search for GO terms, or ortholog *Arabidopsis thaliana* gene IDs. Gene expression graphs can be generated and graphs and data can be downloaded directly from the tool. This tool can be expanded to include new quinoa expression data as it becomes available.

Quinoa Heat Data Explorer was developed using Shiny (http://shiny.rstudio.com/). The code for Quinoa Heat Data Explorer is available at https://github.com/danforthcenter/quinoa-heat-tovar. The complete list of genes in the quinoa genome was associated with their respective *A. thaliana* orthologs available from the Phytozome (Goodstein *et al.*, 2012) quinoa genome annotation (Jarvis *et al.*, 2017). The list of GO terms corresponding to each quinoa gene was obtained by submitting the list of all *A. thaliana* orthologs to Panther (http://pantherdb.org/) (Mi *et al.*, 2013) and retrieving the corresponding GO terms. For each gene, the corresponding corrected differential expression significance value (q value) and the expression value (b value, labelled “de” in Quinoa Heat Data Explorer) as obtained from analysis with Sleuth was added for each treatment and day of heat treatment sampled.

**Figure S1.**
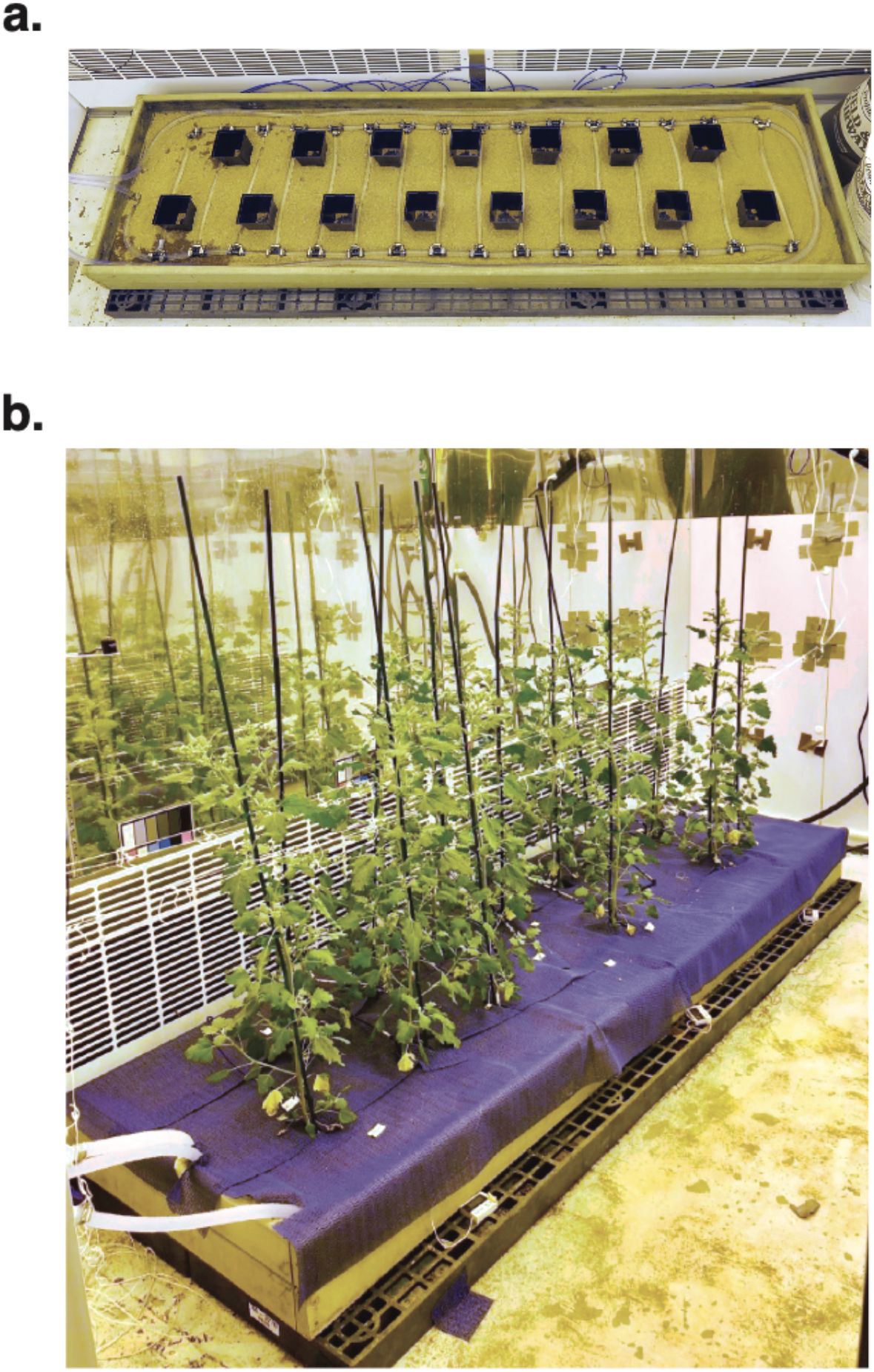
Sandbox system to apply heat and cooling treatments. a) Sandbox system with cooling hose running around pots. b) Sandbox system with quinoa plants during heat treatment.

**Figure S2.**
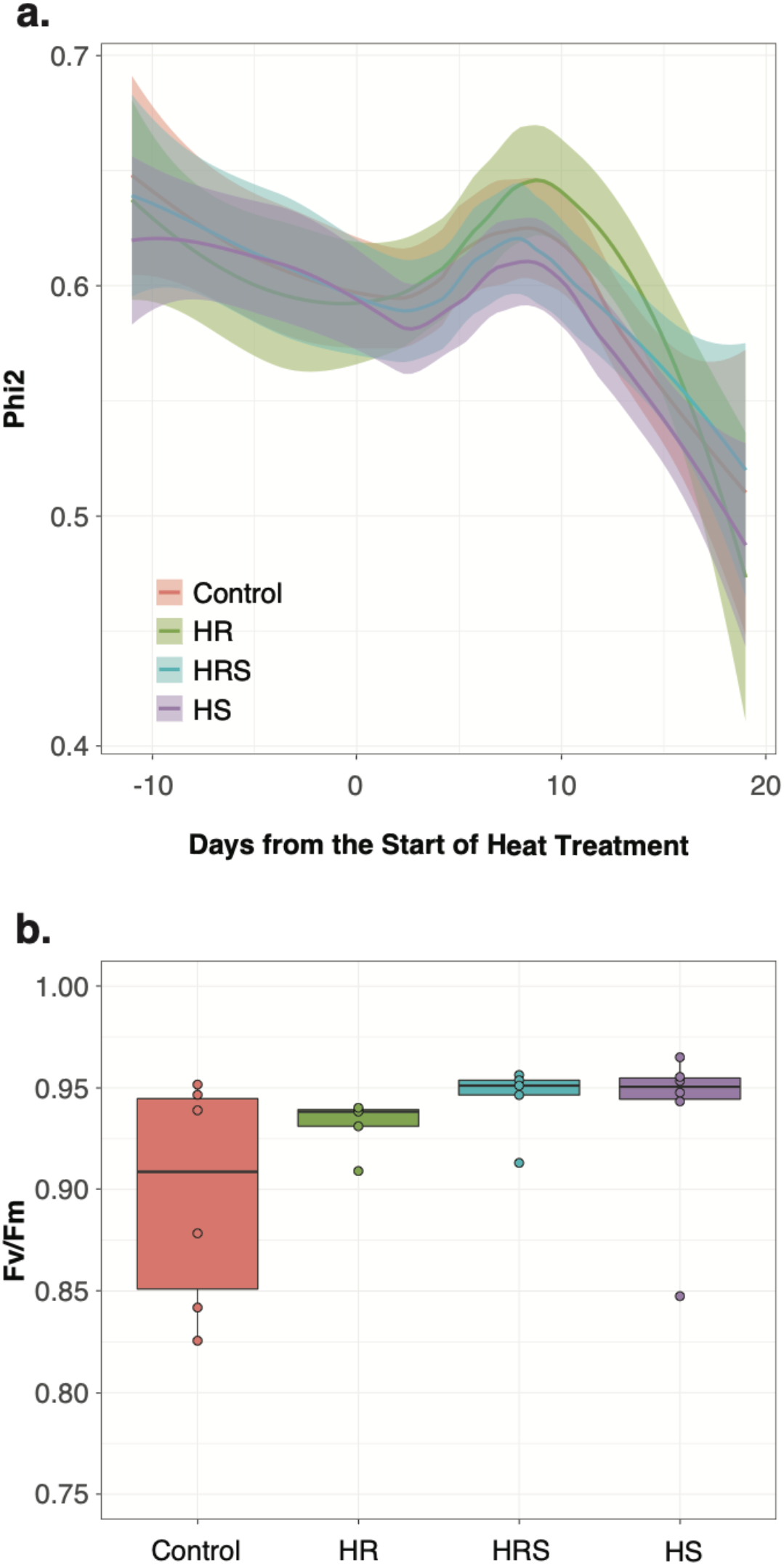
Photosystem II efficiency was not changed by heat treatment. a) Phi2 measured from 10 days before heat treatment started, during heat treatment, and until 8 days after heat treatment ended (n = 3 to 8 plants per timepoint and per treatment). Curves resulting from a LOESS polynomial regression are shown. b) Fv/Fm measured after 11 days in heat treatment per plant for each treatment (n =5 for control and HS; n=6 for HR and HRS).

**Figure S3.**
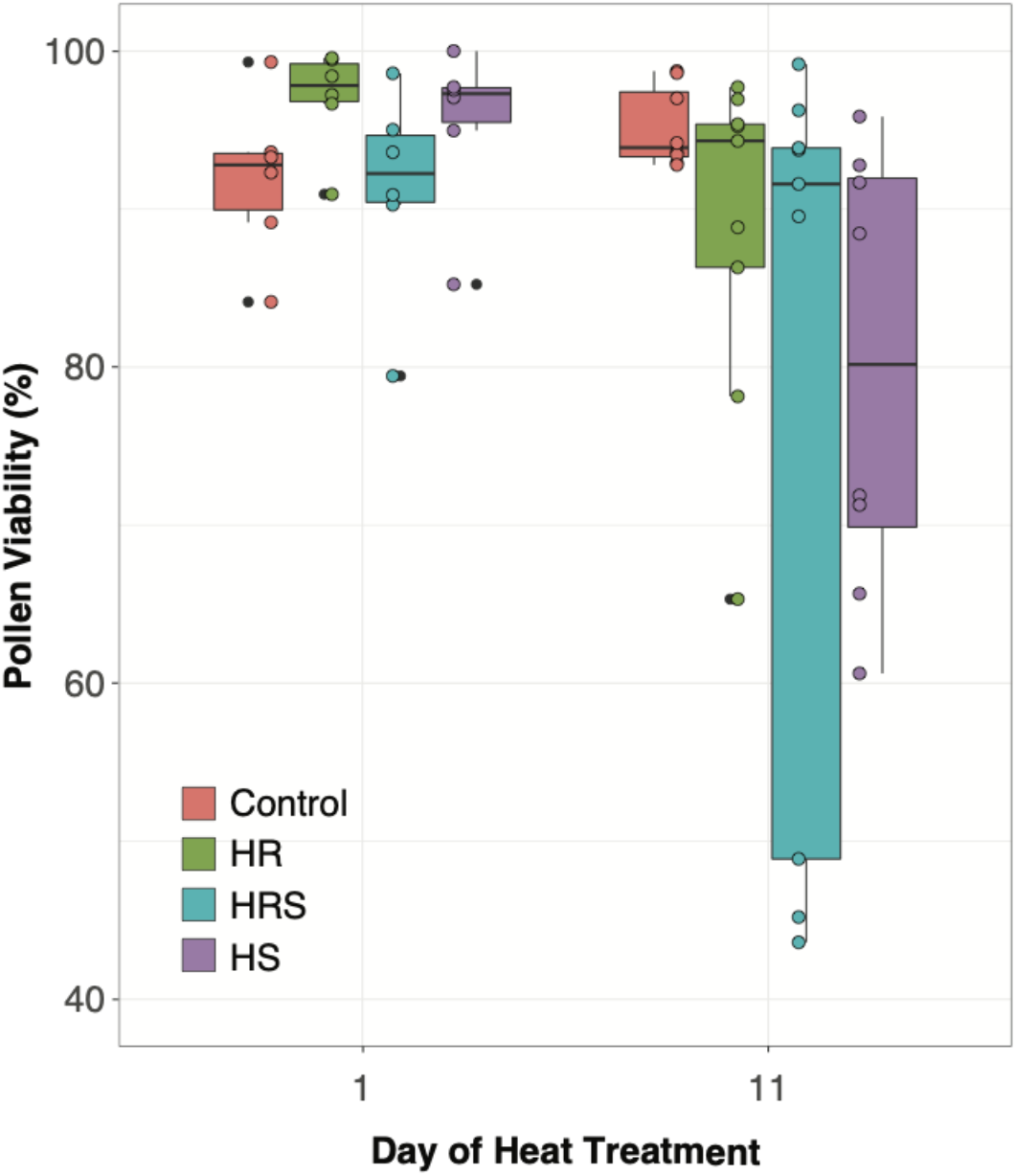
Pollen viability measured during 1 and 11 days of heat treatment (n=6 plants per treatment for day 1, and n=8 plants per treatment for day 11).

**Figure S4.**
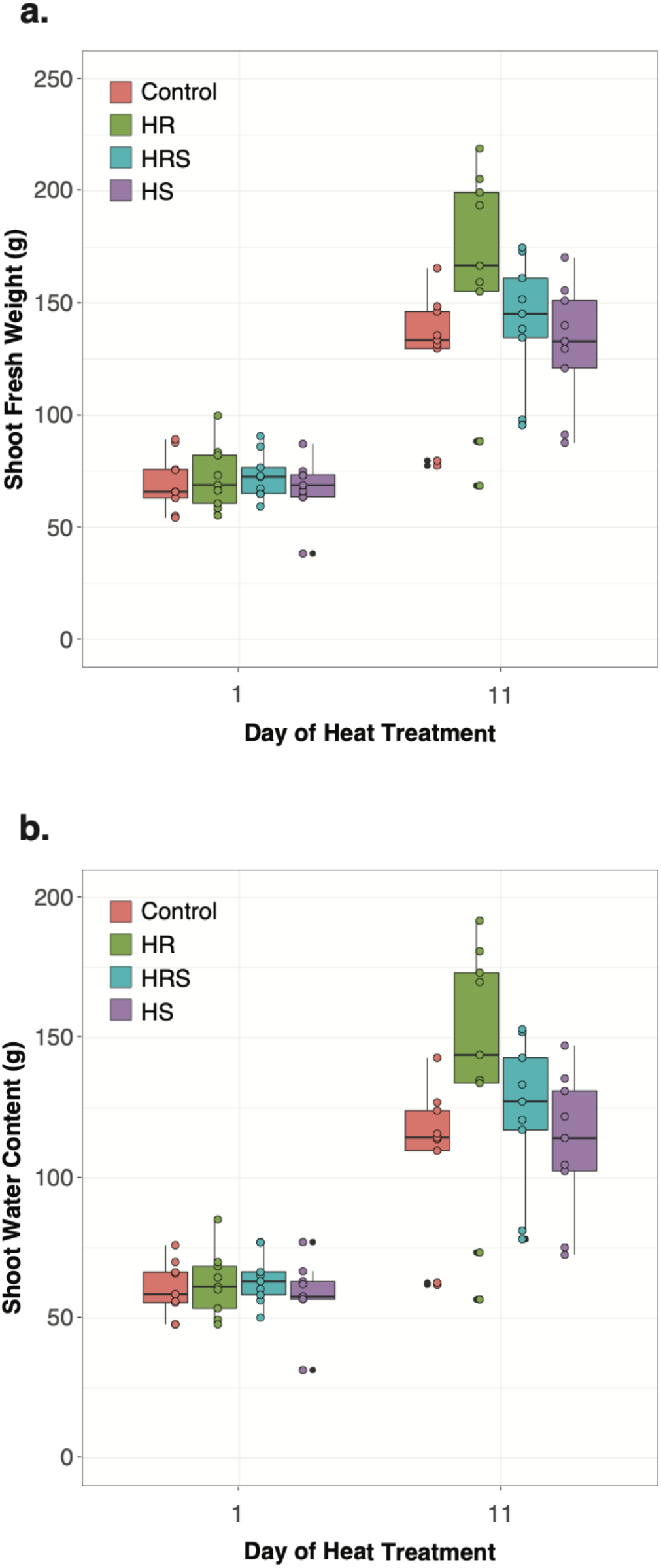
Shoot fresh weight and water content were not modified after heat treatment. a) Shoot fresh weight measured from each plant and for each treatment (n=9). b) Shoot water content measured from each plant and for each treatment (n=9).

**Figure S5.**
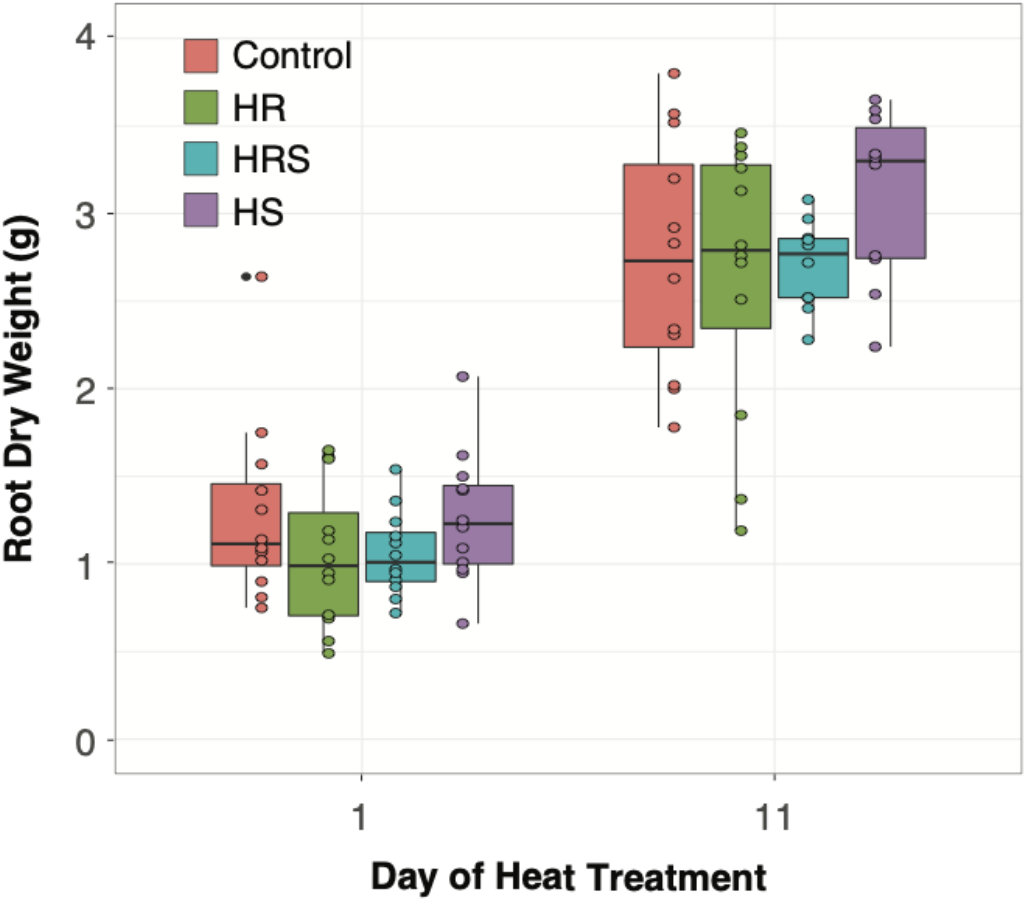
Root dry weight was not significantly affected by heat treatment. Root dry weight measured for each plant and for each treatment at 1 and 11 days of heat treatment (n=129).

## Notes

**Conflict of interest statement** The authors declare no conflict of interest.

https://github.com/danforthcenter/quinoa-heat-tovar

https://doi.org/10.5281/zenodo.3352281

